# Hierarchical dendrite-inspired organic bioelectronic interfaces for neuronal integration

**DOI:** 10.64898/2026.07.30.741673

**Authors:** Kevin Lengefeld, Esther Matamoros, Colin Fernandes, Francesca D’Elia, Giuseppina Iachetta, Elke Brauweiler-Reuters, Nevena Stajković, Michele Dipalo, Claudia Latte Bovio, Andreas Offenhäusser, Simon Musall, Francesco De Angelis, Valeria Criscuolo, Francesca Santoro

## Abstract

Brain–machine interfaces rely on intimate electrical communication between living neurons and artificial materials, yet conventional electrode architectures remain structurally different to neuronal tissue, limiting stable cell–electrode coupling and long-term recording. Although conducting polymers and microstructured interfaces have individually improved the electrochemical and biological properties of neural electrodes, integrating neuromimetic architecture with electrically active organic materials to direct neuron–electrode interactions has remained challenging.

Here we engineer dendrite-inspired organic microelectrodes by programming the self-assembly of conducting polymer fibers directly on microelectrode arrays. By controlling the electrodeposition process, we generate hierarchical architectures that emulate key stages of dendritic development while preserving the relevant electrochemical properties of conducting polymers. These bioinspired interfaces promote nanoscale membrane conformability, reduce the neuron–electrode cleft, and induce localized membrane engulfment of the electrode structures. The enhanced structural integration is accompanied by increased synaptic protein expression and neuronal excitability within mature primary cortical networks. Furthermore, the dendritic electrodes enable localized laser-assisted optoporation, providing transient intracellular electrophysiological access while maintaining extracellular recording functionality.

## Introduction

Neural interfaces have seen significant advancements in recent decades, overcoming the limitations of conventional electrodes, which are typically based on rigid metals or silicon, that exhibit a pronounced mechanical and structural mismatch with the soft and dynamically changing brain environment. This has been shown to lead to foreign body response and the formation of glial scar tissue, which progressively isolates the device and consequently reduces signal fidelity^1,2^. Even when an electrode retains its electrical functionality, the lack of a stable, intimate coupling at the tissue-electrode interface remains a central challenge for chronic applications. Consequently, efforts are being made to improve the integration with neural tissues. Conventional electrode materials such as gold, platinum, and iridium oxide remain the benchmark for electrochemical stability and charge-injection capacity^3^, yet their rigid, planar geometries fall short of addressing this underlying mechanical and structural mismatch.

Conductive polymers have emerged as an alternative material for neural interfaces^4^. Their mixed ionic and electronic conduction and a comparatively large effective surface area result in a reduction of the electrodes’ impedance and an enhancement of charge transfer, resulting in higher signal recordings^5^. In addition, the chemical tunability of these materials enables surface functionalization, thereby facilitating the active regulation of neuronal adhesion, differentiation, and neurite extension^6^. Furthermore, combining conducting polymers with hydrogels reduces electrode stiffness towards the Young’s modulus of neuronal tissue, potentially enabling seamless integration through improved mechanical compliance^7^.

Recently, three-dimensional micro- and nanostructured electrodes have been developed to better match the structural complexity of neural tissue. Gold mushroom-shaped microelectrodes, for instance, promote tight, engulfment-like membrane wrapping that enables multi-site, intracellular-like recordings from individual neurons^8^. Porous and hierarchical surface features have also been shown to strengthen functional connectivity across neuronal populations^9^ and to reshape network architecture toward small-world topologies^10^, indicating that topographical engineering can influence tissue integration at both the cellular and network scale.

In this context, neuromorphic interfaces offer a conceptual approach that goes beyond traditional electrode architectures by taking inspiration from the structural and organizational principles of neural systems^11^. This approach emulates the structural and mechanical cues presented by neurons themselves, while also incorporating electrical, ionic, and chemical modalities as complementary dimensions^12^. This structural emulation, together with the accompanying functional integration, supports closer bidirectional communication between the brain and the device. Micro- and nanostructures resembling the structure of dendritic spines have been found to influence the early outgrowth of neural cells^13^. A related strategy exploits structures that mimic neurites rather than spines. Aligned, eumelanin-coated electrospun PLA microfibers have been used to guide neuronal spreading, alignment, and maturation, directly linking neurite-like topographical cues to improved cell-material interaction. Moreover, the use of neurite-like electrodes, matching both the size and mechanical properties of neurons, has enabled stable long-term recording of electrical brain activity while also reducing the associated inflammatory response^14^.

Despite significant advances in bioelectronic interfaces, current neuron–electrode platforms largely optimize either electrical performance or structural biomimicry but rarely integrate both to actively direct neuron–electrode interactions and establish functional neurohybrid networks. An interface that combines neuromimetic morphology with electrically active organic materials to control neuronal organization at the nanoscale, promote intimate neuron–electrode coupling, modulate synaptic connectivity and network activity, and enable intracellular electrophysiological access has remained elusive.

Here, we engineer neuromimetic PEDOT-based organic microelectrodes with dendrite-inspired architectures on microelectrode arrays (MEAs) that structurally emulate key features of neuronal processes. By guiding neuronal integration at the nanoscale, these bioinspired electrodes establish an exceptionally intimate neuron–electrode interface with primary cortical neurons, resulting in enhanced physical coupling between neuronal membranes and the electrode surface. This close integration is accompanied by a trend towards increased expression of synaptic proteins at the electrode interface, indicating possible synaptic organization on the neuromimetic structures. Electrophysiological recordings further reveal enhanced neuronal excitability and increased functional connectivity in mature neuronal networks cultured on the dendrite-inspired electrodes, demonstrating that the engineered morphology actively influences network-level electrical dynamics. Finally, we show that the same neuromimetic electrodes enable localized laser-assisted optoporation, providing transient intracellular access for electrophysiological sensing while preserving the extracellular recording capability of the platform (**Fig. 1a**). Together, these results establish a multifunctional neurohybrid interface that couples structural biomimicry with organic bioelectronics to control neuron-electrode interactions while enabling both extracellular modulation and intracellular interrogation.

## Results

### Hierarchical electrode topography promotes intimate neuron–electrode coupling

**Figure 1.**
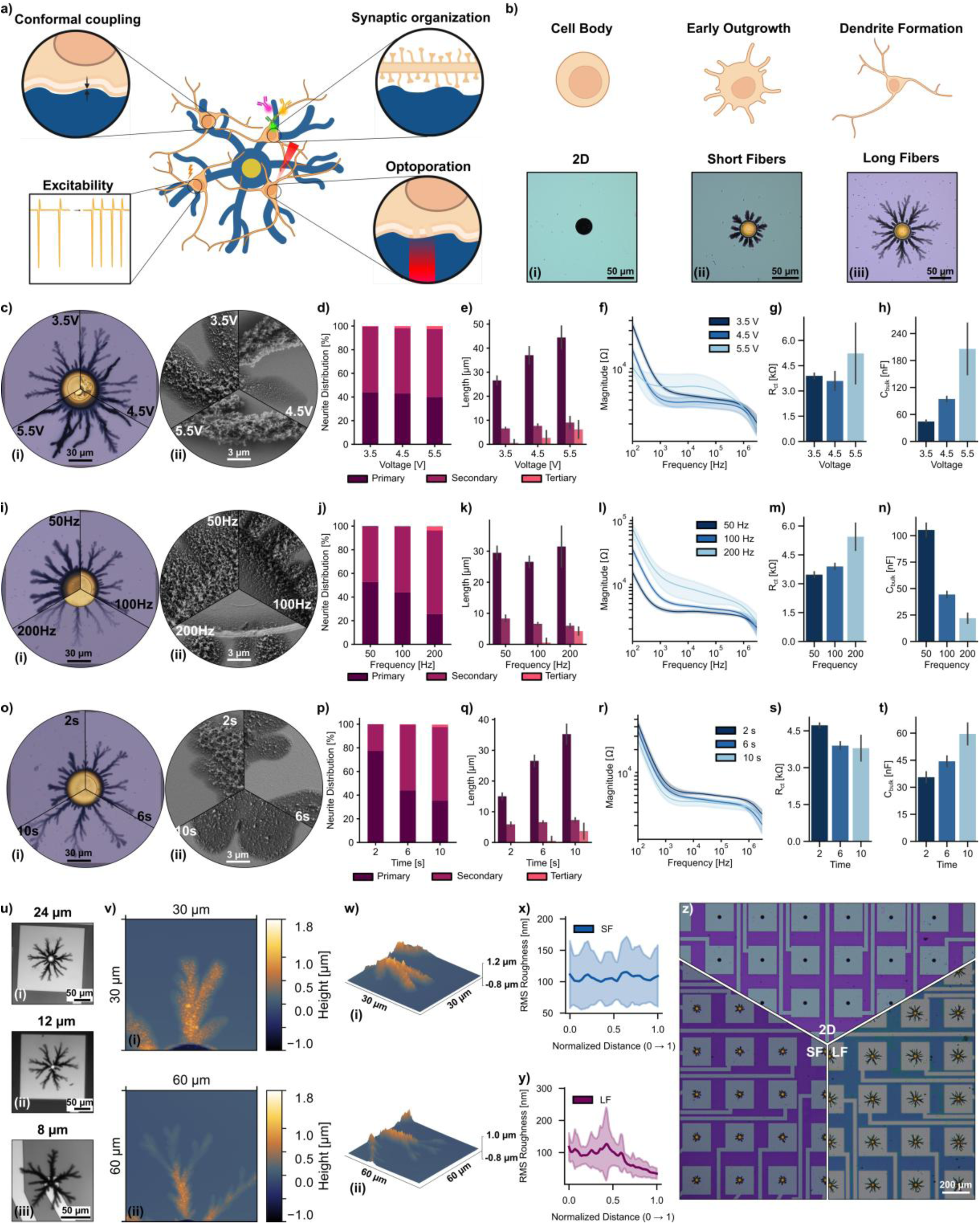
Engineering and electrochemical characterization of neuromimetic PEDOT:PF_6_ microelectrodes. **a**, Schematic illustrating the concept and key findings of the neuromimetic electrode platform. **b,** Optical micrograph of PEDOT:PF_6_ electrodes, namely 2D (**i**), SF (**ii**) and LF (**iii**), engineered to mimic neuronal outgrowth stages. 2D electrodes were deposited galvanostatically (69.1 nA, 8 s), SF and LF electrodes were deposited applying dynamic biphasic pulses (±3.5 V, 100 Hz) for 2 and 6 s, respectively. **c**-**h**, Variation of the deposition voltage (3.5, 4.5 and 5.5 V; deposition frequency, 100 Hz, deposition time, 6 s): representative optical (**c**,**i**) and scanning electron micrographs (**c**,**ii**), quantification of the branch (neurite-like) distribution (**d**) and neurite length (**e**), impedance magnitude spectra (**f**), fitted charge transfer resistance (**g**) and bulk capacitance (**h**). **i-n**, Variation of the deposition frequency (50, 100, 200 Hz; deposition voltage, 3.5 V; deposition time, 6 s): representative optical (**i**,**i**) and scanning electron micrographs (**i**,**ii**) quantification of the branch distribution (**j**) and neurite length (**k**), impedance magnitude spectra (**l**), fitted charge transfer resistance (**m**) and bulk capacitance (**n**). **o**-**t**, Variation of the deposition time (2, 6, 10 s; deposition voltage, 3.5 V, deposition frequency, 100 Hz): representative optical (**o**,**i**) and scanning electron micrographs (**o**,**ii**), quantification of the branch distribution (**p**) and neurite length (**q**), impedance magnitude spectra (**r**), fitted charge transfer resistance (**s**) and bulk capacitance (**t**). **u**, Neuromimetic fiber electrodes scaled to 24 (**i**), 12 (**ii**) and 8 µm (**iii**) electrodes. **v**-**y**, AFM analysis of the fiber topography: representative AFM scans of SF (**u**,**i**) and LF (**u**,**ii**) branches, three-dimensional reconstruction of representative SF (**w**,**i**) and LF (**w**,**ii**) branches, RMS roughness along the SF (**x**) and LF (**y**) branches, normalized to the maximum length of the SF or LF branches. **z**, Large-area optical micrographs of 2D, SF and LF electrodes integrated on MEAs.

To promote structural integration between living neurons and electronic interfaces, we engineered neuromimetic PEDOT:PF_6_ microelectrodes that recapitulate key stages of dendritic development. Specifically, planar (2D), short-fiber (SF) and long-fiber (LF) electrodes were designed to resemble the neuronal soma, early neurite outgrowth and mature dendritic arborization, respectively (**Fig. 1b**). The electrodes were fabricated using an electrodeposition strategy in which biphasic voltage pulses drive the self-organized growth of dendritic PEDOT fibers^15^ directly from individual MEA electrodes towards a Pt counter electrode immersed in the monomer solution (**Supplementary Fig. 1**). By tuning the deposition conditions, this approach enables programmable control over the morphology of the resulting electrodes, providing a versatile platform for engineering neuromimetic bioelectronic interfaces. To establish design rules governing electrode morphology, we systematically varied the deposition voltage, pulse duration, *i.e*., signal frequency, and deposition time using 50 μm-diameter electrodes. Deposition voltages ranged from 3.5 to 5.5 V, pulse frequencies from 50 to 200 Hz, and deposition times from 2 to 10 s. Representative morphologies obtained by varying the deposition voltage are shown in **Fig. 1c**. Increasing the deposition voltage promoted the formation of progressively more branched dendritic architectures, although partial delamination of the branches from the passivation layer was observed at higher voltages (**Fig. 1c-ii**), indicating a trade-off between structural complexity and mechanical stability. To quantitatively assess the resulting architectures, we adapted a neurite tracing workflow previously used for neuronal morphometric analysis (Material and Methods) to the dendritic PEDOT structures (**Supplementary Fig. 2**). This analysis enabled direct quantification of branch distribution and branch length, providing a framework for comparing the neuromimetic morphologies generated under different deposition conditions. Quantitative morphometric analysis revealed that the branching hierarchy was largely preserved across the investigated deposition voltages (**Fig. 1d**), whereas branch length increased with increasing voltage (**Fig. 1e**), consistent with an enhanced polymerization rate at higher applied potentials^16^.

Electrochemical impedance spectroscopy (EIS) (**Fig. 1f**) further demonstrated that electrode morphology directly governs the electrochemical properties of the interface. Increasing the deposition voltage initially reduced the charge transfer resistance before increasing at 5.5 V (**Fig. 1g**), indicating that excessive growth compromises the effective electrochemically accessible surface. Moreover, the larger variability observed at 5.5 V suggests reduced reproducibility under highly accelerated growth conditions. On the other hand, increasing the deposition voltage resulted in a larger bulk capacitance (**Fig, 1h**), reflecting the increased amount of deposited material.^17^

Furthermore, signal frequency provided an independent strategy to tune the electrode’s architecture (**Fig. 1i**). In fact, increasing the signal frequency progressively reduced the amount of deposited material while promoting the formation of higher order branches^18^ (**Fig. 1j**) without substantially affecting branch length (**Fig. 1k**). This morphological transition was accompanied by a change in the electrochemical properties (**Fig. 1l**). An increase in charge-transfer resistance (**Fig. 1m**) and a decrease in bulk capacitance (**Fig. 1n**), consistent with reduced polymer deposition at shorter pulse durations. We attribute this behavior to the increasing contribution of electrical double-layer charging, which limits the faradaic polymerization process at high frequencies and consequently defines an upper frequency limit for efficient fiber growth.

In contrast, extending the deposition time resulted in thinner branch ends, suggesting a reduced polymerization rate (**Fig. 1o**). Simultaneously, longer deposition times promoted the formation of higher order branches (**Fig, 1p**) resulting in progressively more complex dendritic architectures, suggesting that the growth is initially diffusion-limited as the close proximity of polymerization sites in the early growth restricts the local transport of monomers and counterions. As anticipated, an increase in the deposition time resulted in the elongation of the branches (**Fig. 1q**). Accordingly, a continuous decrease in the impedance can be observed (**Fig. 1r**), as a consequence of the reduced charge-transfer resistance (**Fig. 1s**) and increased bulk capacitance (**Fig. 1t**), consistent with the formation of larger electrochemically active volumes. Notably, the charge-transfer resistance reached a plateau beyond 6 seconds despite continued increases in branch length and deposited material, indicating diminishing electrochemical gains with prolonged deposition.

Together, these results establish deposition voltage, pulse frequency and deposition time as orthogonal parameters for programming the morphology and electrochemical properties of neuromimetic PEDOT:PF6 electrodes. Based on the balance between structural complexity, mechanical stability and electrochemical reproducibility, a deposition voltage of 3.5 V and a signal frequency of 100 Hz were selected for subsequent studies. To further approximate the dimensions of neuronal soma, the deposition strategy was translated to electrodes with diameters of 24, 12 and 8 μm. To preserve the characteristic dendritic morphology during miniaturization, the deposition voltage was scaled with electrode diameter and gradually increased in 8 mV increments during growth until reaching 3.5 V (**Supplementary Fig. 3**). This strategy yielded well-defined neuromimetic fiber architectures across all electrode sizes (**Fig. 1u**), demonstrating the scalability of the fabrication.

Based on the optimization study, three electrode architectures were selected for subsequent experiments: 2D, SF, and LF, representing successive stages of neuronal process development. Compared with uncoated gold electrodes, all PEDOT:PF_6_ electrodes exhibited impedances that were several orders of magnitude lower across the biologically relevant frequency range^19^ (**Supplementary Fig. 4a,b**). This was accompanied by a reduced charge-transfer resistance (**Supplementary Fig. 4c**), indicative of an increased electrochemically accessible surface area, and a substantially higher bulk capacitance arising from the volumetric charge storage of PEDOT:PF_6_ (**Supplementary Fig. 4d**). The enhanced capacitive behavior was further confirmed by cyclic voltammetry (**Supplementary Fig. 4e**).

Atomic force microscopy (AFM) revealed pronounced differences in nanoscale topography between the three designs (**Fig. 1v,w**). Whereas the SF electrodes exhibited a nearly constant root-mean-square (RMS) roughness (**Fig. 1x**) of approximately 100 nm along the branches, the LF electrodes showed a progressive decrease in roughness towards the branch tips (**Fig. 1y** and **Supplementary Fig. 5a,b**), consistent with branch thinning during growth. In contrast, uncoated gold and planar PEDOT:PF_6_ electrodes exhibited RMS roughness values of only 2-3 nm and approximately 38 nm, respectively (**Supplementary Fig. 5c,d**). The fiber electrodes therefore combine hierarchical dendritic architectures with nanoscale surface features that more closely resemble the structural complexity of neuronal processes than conventional planar electrodes and were subsequently integrated across complete microelectrode arrays (**Fig. 1z**).

**Figure 2.**
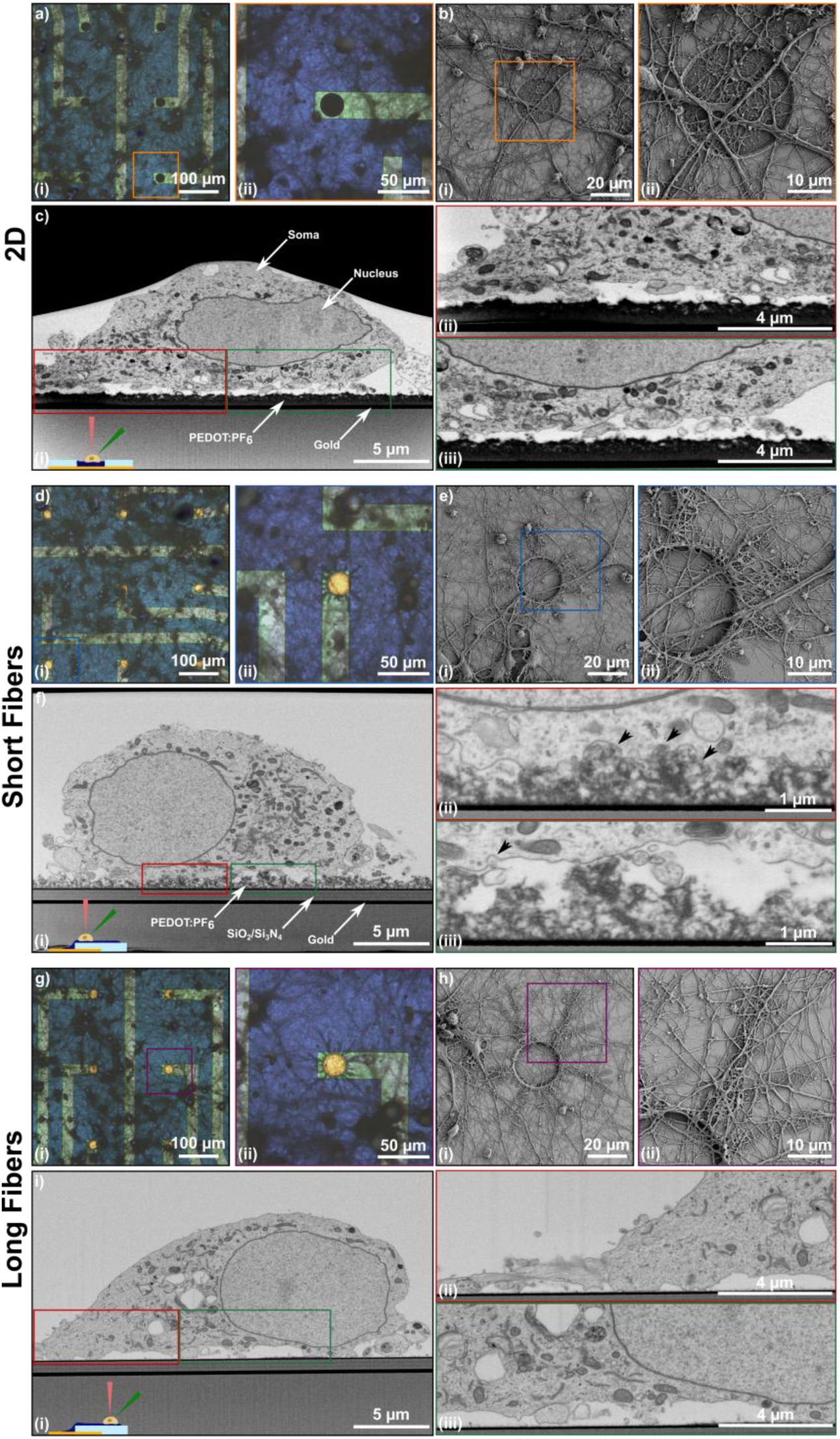
Hierarchical electrode topography promotes intimate neuron–electrode coupling. **a**, Optical micrograph of resin embedded primary cortical neurons on a 2D microelectrode array (**i**) and single electrode (**ii**). **b**, Representative scanning electron micrographs of a neuron interfacing a 2D electrode (**i**) and higher magnification view (**ii**). **c**, FIB-SEM micrograph cross-section of the neuron-electrode interface on a 2D electrode (**i**), showing the membrane-electrode cleft (**ii**,**iii**). **d**, Optical micrograph of resin embedded neurons on a SF microelectrode array (**i**) and single electrode (**ii**). **e**, Representative scanning electron micrograph of a neuron at interfacing a SF electrode (**i**) showing neurite extension along individual fiber branches (**ii**). **f**, Representative FIB-SEM micrograph cross-section acquired at the base of a LF electrode (**i**), illustrating the intimate membrane conformability (**ii**, **iii**) observed at both SF branches and the base of LF branches. **g**, Optical micrograph of resin embedded neurons at DIV14 on a LF microelectrode array (**i**) and single electrode (**ii**). **h**, Representative SEM micrograph of neurons at DIV5 on a LF electrode (**i**) showing neurite extension along the LF branches (**ii**). **i**, FIB-SEM cross-section at the tip of an LF branch (**i**), revealing reduced membrane conformability compared with the base (**ii**, **iii**).

Primary cortical neurons were cultured on 2D, SF, and LF microelectrode arrays to evaluate the biological compatibility of the neuromimetic interfaces. Biocompatibility assays confirmed cell viability comparable to the glass control on all electrode architectures, demonstrating that the conducting polymer coatings support long-term neuronal culture (**Supplementary Fig. 6**). To determine how the engineered topography influences neuron-electrode interactions, cultures were analyzed by optical microscopy, scanning electron microscopy (SEM), and focused ion beam (FIB)-SEM cross-sectioning (**Fig. 2**). FIB-SEM cross-sections were acquired at 5 and 7 days *in vitro* (DIV5 and DIV7, respectively), to avoid dense neurite coverage. On 2D electrodes, neurons covered the recording sites and extended neurites across the PEDOT:PF_6_ surface (**Fig. 2a,b**). However, FIB-SEM cross-sections revealed a pronounced membrane-electrode cleft beneath the neuronal soma, indicative of limited physical coupling to the planar interface (**Fig. 2c**).

In contrast, neurons cultured on SF electrodes established a more intimate interface: neurites frequently extended along individual fiber branches (**Fig. 2d,e**), whereas FIB-SEM cross-sections revealed that the plasma membrane closely conformed to the underlying dendritic topography, substantially reducing the membrane-electrode cleft (**Fig. 2f**). Membrane invaginations surrounding individual fiber branches were consistently observed, suggesting active membrane engulfment of the structures, a coupling mechanism previously reported for micro- and nanostructured biointerfaces^20^. Similar conformal membrane wrapping was also observed where neurites interfaced with the fiber branches (**Supplementary Fig. 7**), indicating that the enhanced coupling is not restricted to the neuronal soma.

LF electrodes exhibited the same overall integration behavior (**Fig. 2g,h**). However, cross-sections acquired near the distal branch tips revealed a less conformal interface than at the branch base (**Fig. 2i**). This reduced membrane wrapping correlates with the progressive decrease in branch diameter and nanoscale roughness towards the fiber tip, suggesting that local geometrical features govern the strength of neuron-electrode coupling.

Together, these observations demonstrate that hierarchical dendritic topography promotes nanoscale membrane conformability and intimate neuron-electrode integration, providing a structural basis for the synaptic organization and electrophysiological coupling described below.

**Figure 3.**
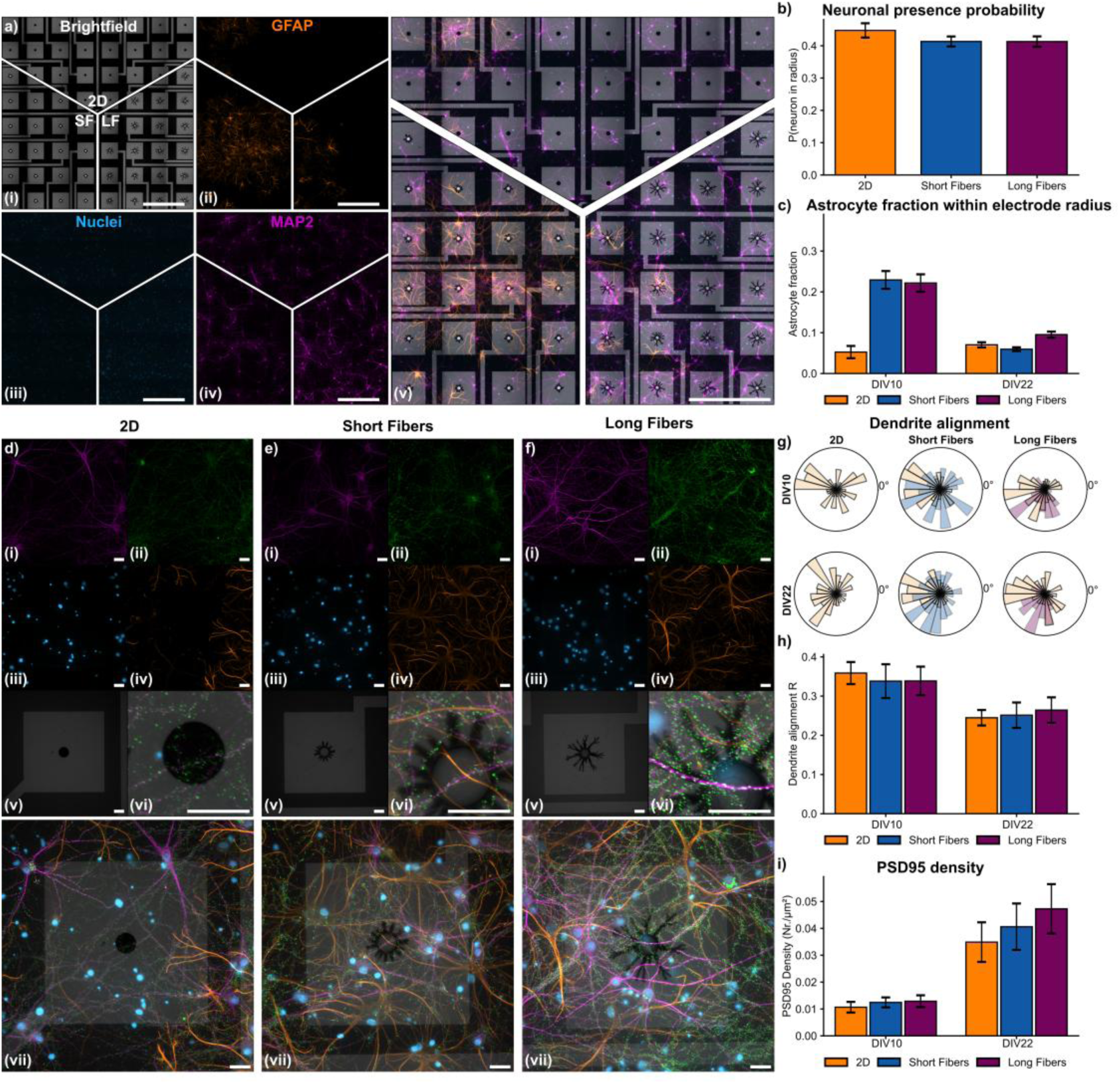
Neuromimetic electrodes promote localized synaptic organization. **a**, Representative tiled optical micrographs of primary cortical neurons cultured on 2D, SF and LF microelectrode arrays at DIV22. Micrographs depict the electrode array (brightfield; **i**), astrocytes (GFAP; **ii**), nuclei (DAPI; **iii**), dendrites (MAP2; **iv**) and an overlay of all channels (**v**). Scale bars denote 500 µm. **b**, Probability of a neuronal soma within a radius of 80 µm of the electrode’s centre for DIV10 and DIV22 combined. **c**, Fraction of astrocytes within the same radius at DIV10 and DIV22. **d**-**f**, Representative single electrode optical micrographs of primary cortical neurons at DIV22 on a 2D (**d**), SF (**e**) and LF (**f**) electrode, depicting dendrites (MAP2; **i**), excitatory postsynaptic sites (PSD95; **ii**), nuclei (DAPI; **iii**), astrocytes (GFAP; **iv**), electrode (brightfield; **v**) and an overlay of all channels of the electrodes region of interest (ROI; **vi**) and full field of view (**vii**). Scale bars denote 20 µm. **g**, Radial distribution of neurite orientation relative to the electrode for 2D, SF and LF electrode at DIV10 and DIV22. **h**, Alignment coefficient quantifying neurite orientation at DIV10 and DIV22. **i**, Density of PSD95 on the electrode surface at DIV10 and DIV22. A trend of increased PSD95 accumulation on SF and LF electrodes, together with the absence of preferential neurite alignment, suggests a possible synaptic organization without altering the global architecture of the neuronal network. Brightness and contrast of fluorescence micrographs were adjusted linearly and identically across within the same type. Data represent n = 3 per condition from N = 3 independent experiments for each DIV. Graphs represent the mean ± sem.

### Neuromimetic electrodes promote synaptic organization

To determine whether the engineered electrode architectures influence neuronal organization beyond the electrode interface, primary cortical cultures were immunostained at DIV10 and DIV22, to investigate possible structural changes induced at different neuronal maturation stages. Fluorescence micrographs were acquired for dendrites (microtubule-associated protein 2; MAP2), astrocytes (glial fibrillary acidic protein; GFAP), nuclei (4′,6-diamidino-2-phenylindole; DAPI) and the electrodes structure (brightfield) (**Fig. 3a**). Quantification of the probability of neuronal soma being within a 80 μm radius from the electrodes center of 2D, SF and LF electrodes revealed no difference (**Fig. 3b**), indicating an absence of neuronal clustering in the electrodes surrounding. Likewise, a slight but not significant increase in the fraction of astrocytes in the same radius for SF and LF electrodes was visible at DIV10, which is reduced at DIV22, suggesting that the fiber structures do not trigger an increased astrocyte coverage (**Fig. 3c**). Together, these observations indicate that the dendrite-inspired interfaces do not alter the spatial distribution or behavior of neuronal cells. To further investigate the local neuron–electrode interface, higher-magnification optical micrographs of individual electrodes were acquired, including immunostaining for excitatory postsynaptic sites (postsynaptic density protein 95; PSD95) (**Fig. 3d–f** and **Supplementary Fig. 8a-c**). PSD95 was chosen as synaptic marker due to its central role in anchoring neurotransmitter receptors and driving the maturation of excitatory synapses. Neurite orientation relative to the dendritic electrode geometry was first quantified to determine whether the fiber architectures act as contact-guidance cues. Radial orientation analysis revealed no preferential alignment of neurites with the direction of the electrode branches (**Fig. 3g**), and the corresponding alignment coefficient remained comparable across electrode architectures and developmental stages (**Fig. 3h**). These findings indicate that, despite their biomimetic morphology, the dendritic electrodes do not impose long-range neurite guidance or anisotropic growth on the surrounding neuronal network. Instead, their influence appears to be confined to the electrode interface, where neurons establish direct physical contact with the hierarchical topography.

In contrast to the absence of large-scale structural reorganization, possible differences emerged at the molecular level: PSD95 density increased from DIV10 to DIV22 on all electrode architectures, consistent with progressive synaptic maturation of the cultures^21^ (**Fig. 3i**). Notably, both SF and LF electrodes exhibited a trend towards a greater accumulation of PSD95 than 2D electrodes, indicating that hierarchical dendritic topography might promote synaptic organization at the neuron-electrode interface. Together with the ultrastructural evidence of enhanced membrane conformability, these results suggest that neuromimetic architectures strengthens structural integration with neurons without disrupting the overall organization of the surrounding network.

**Figure 4.**
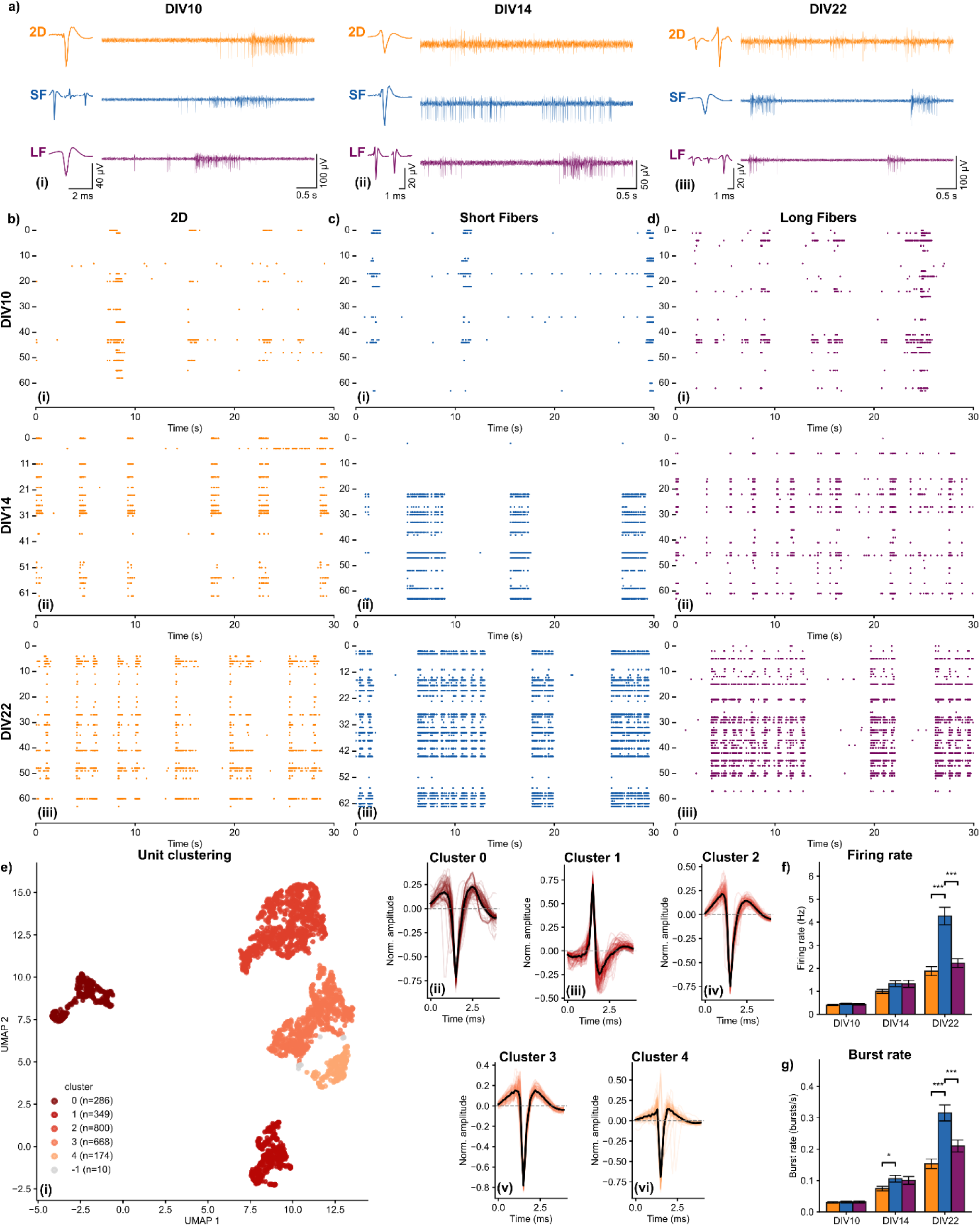
Dendrite-inspired electrodes enhance neuronal network excitability. **a**, Representative extracellular traces from primary cortical neurons cultured on 2D, SF and LF microelectrode arrays at DIV10 (**i**), DIV14 (**ii**) and DIV22 (**iii**). Detected units are depicted on the left side of the corresponding trace. **b-d**, Representative raster plots on 2D (**b**), SF (**c**) and LF (**d**) MEAs at DIV10 (**i**), DIV14 (**ii**) and DIV22 (**iii**), illustrating the increased activity throughout the maturation of the neuronal network. **e**, Unsupervised clustering of detected and curated extracellular units using UMAP and HDBSCAN (**i**), overlayed unit waveforms and mean waveforms of each cluster (**ii**-**vi**). **f**,**g**, Quantification of firing rate (**f**) and burst rate (**g**) across the DIVs and electrode types. Data represent n = 3 per condition from N = 3 independent experiments for each DIV. Graphs represent the mean ± sem.

### Neuromimetic electrodes modulate network electrophysiology

To determine whether the enhanced structural integration translates into functional changes at the network level, spontaneous extracellular electrophysiological recordings were performed at DIV10, DIV14 and DIV22. Representative recordings and raster plots revealed the expected maturation of spontaneous network activity over time, characterized by progressively synchronized bursting across the neuronal cultures (**Fig. 4a-d**), consistent with the development of functional cortical networks *in vitro*^22,23^.

Detected extracellular units were isolated by spike sorting and filtered according to established quality metrics^24^ to identify single unit activity (Materials and Methods). Unsupervised clustering of waveform features using uniform manifold approximation and projection (UMAP) and hierarchical density-based spatial clustering of applications with noise (HDBSCAN) resolved five distinct unit classes that were consistently identified across electrode architectures (**Fig. 4e**), indicating that differences in neuronal activity are unlikely to arise from systematic biases in spike classification (**Supplementary Figure 9f**). Additional quality metrics and representative autocorrelograms are provided in **Supplementary Fig. 9g.**

Quantitative analysis revealed a progressive increase in both firing rate and burst rate with neuronal maturation for all electrode architectures (**Fig. 4f,g**). Notably, neurons interfaced with SF electrodes consistently exhibited significantly higher firing and burst rates than those cultured on 2D or LF electrodes. This increase was coupled to longer burst durations and a greater number of spikes per burst at DIV22 (**Supplementary Fig. 10c,d),** demonstrating that the dendrite-inspired topography not only supports electrical recording but actively enhances neuronal excitability. Together with the increased membrane conformability and localized enrichment of PSD95 observed above, these findings suggest that intimate structural coupling at the neuron-electrode interface promotes more effective functional integration between living neurons and the bioelectronic substrate.

In contrast, LF electrodes did not exhibit a comparable increase in firing activity relative to 2D electrodes. We attribute this primarily to the different coupling mechanisms along the LF branch. As mentioned previously, the gradual reduction in branch diameter and nanoscale roughness (**Figure 1y**) towards the fiber tip is expected to diminish membrane conformability (**Fig. 2i**), attenuating possible local topographical effects that enhance neuronal excitability. Collectively, these observations identify branch topography as a critical determinant of both the biological and electrical performance of dendrite-inspired bioelectronic interfaces.

**Figure 5.**
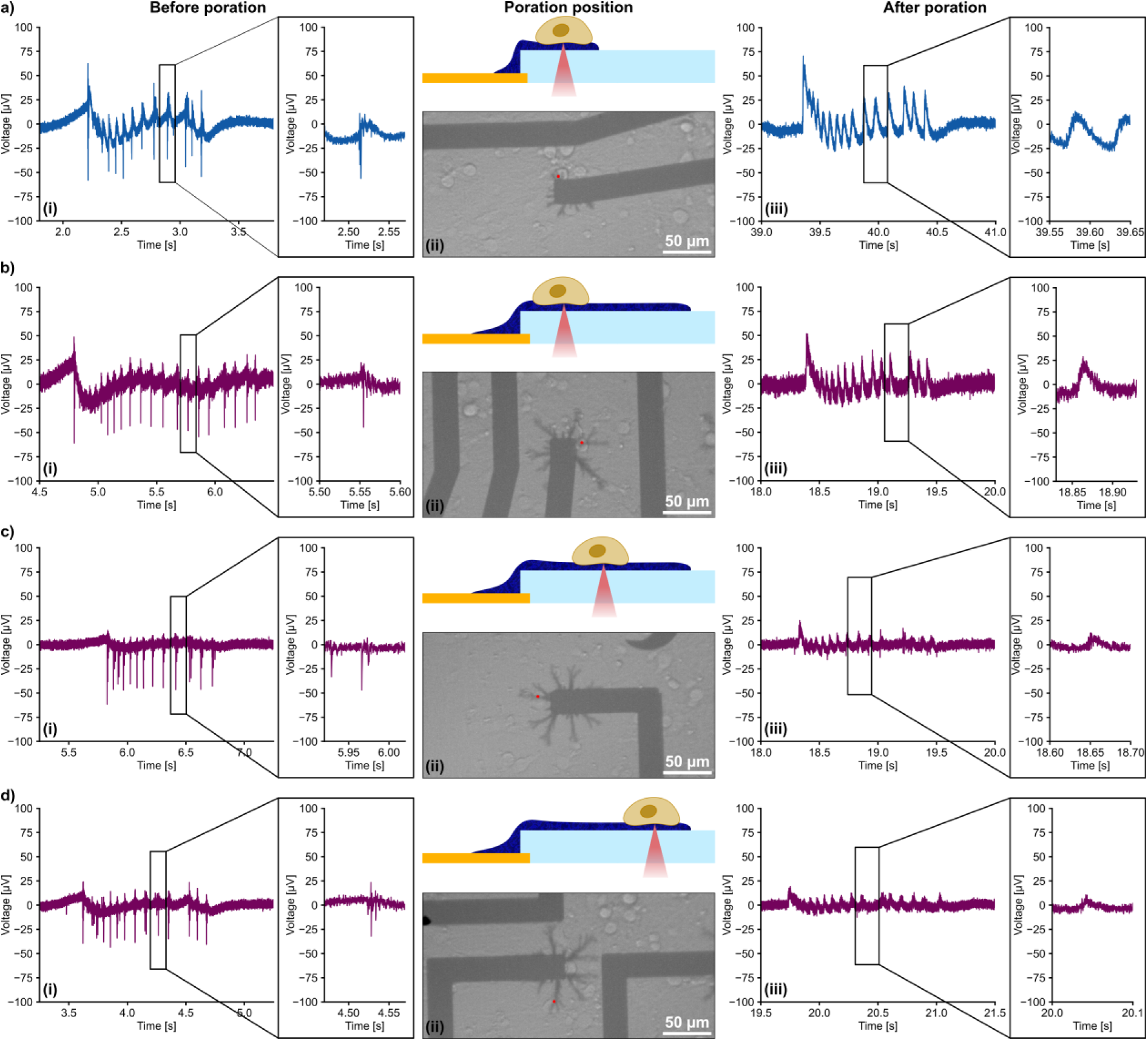
Localized optoporation enables intracellular electrophysiological access through dendrite-inspired electrodes. **a**, Representative optoporation performed on a SF electrode showing the extracellular recording before optoporation (**i**), the laser targeting site on the fiber branch (**ii**), and the in-cell recording following optoporation (**iii**). **b–d,** Representative optoporations performed on LF electrodes localized at the branch base (**b**), intermediate branch position (**c**) and distal branch tip (**d**), each showing the recording before optoporation (**i**), the laser targeting site (**ii**), and the intracellular-like signal following optoporation (**iii**). The progressive reduction in signal quality towards the branch tip reflects the position-dependent neuron-electrode coupling along the dendritic architecture.

### Localized optoporation enables intracellular electrophysiological access

To assess whether the neuromimetic electrodes are compatible with intracellular interfacing, localized laser-assisted optoporation^25,26^ was performed at the neuron–electrode interface. Preliminary experiments were conducted using a 1064 nm laser (spot diameter ∼2 µm) on human induced pluripotent stem cell-derived cardiomyocytes (hiPSC-CMs) cultured on the PEDOT:PF_6_ electrode (**Supplementary Fig. 11**), showing typical intracellular action potential signals obtained for optoporation on cardiomyocytes. Further, to assess the structural organization of hiPSC-CMs on the substrate, cells were immunostained at DIV14 for cardiac troponin T (cTnT), a key protein essential for sarcomeric contraction, and counterstained for nuclei (DAPI). Fluorescence micrographs revealed the formation of a confluent cardiomyocyte layer on top of the LF electrodes, displaying characteristic cTnT distribution with visible striations (**Supplementary Fig. 12**). Regarding the optoporation on neurons, a focused 976 nm laser (spot diameter ∼2 μm) was focused between the neuronal soma and individual fiber branches, transiently permeabilizing the plasma membrane while preserving the extracellular recording configuration.

Representative recordings obtained from SF electrodes are shown in **Fig. 5a**. Before optoporation, neurons exhibited characteristic negative extracellular action potentials (**Fig. 5a-i**). Following laser stimulation at the fiber branch (**Fig. 5a-ii**), the recorded waveform underwent a polarity inversion and evolved into a prolonged positive in-cell^27^ potential that remained temporally aligned with the prior extracellular action potentials (**Fig. 5a-iii**). These recordings indicate successful transient intracellular access through the dendrite-inspired electrode architecture. Notably, due to the fiber architectures extending further than the electrodes opening, the obtained optoporations can be solely attributed to the interaction between the laser and PEDOT:PF6, ultimately allowing to exclude possible effects arising from a metal layer beneath, typically necessary in PEDOT coated electrodes^28^.

To determine whether intracellular access depends on the local electrode morphology, optoporation was performed at different positions along LF electrodes. Optooration at the base of the branches produced responses comparable to those observed on SF electrodes (**Fig. 5b**), whereas recordings obtained progressively closer to the branch tip exhibited a gradual reduction in signal-to-noise ratio (**Fig. 5c,d**). This positional dependence mirrors the reduced electrophysiological performance observed for distal branch regions and is consistent with the progressive decrease in branch thickness and nanoscale roughness towards the fiber tip. For comparison, optoporation on planar electrodes (**Supplementary Fig. 13**) produced intracellular-like waveforms containing superimposed positive action potentials, consistent with previous reports using PEDOT-based microelectrodes^29^. These individual spikes were less pronounced on the fiber electrodes, where they were not readily distinguishable from the background noise. Together, these findings demonstrate that the dendrite-inspired electrodes combine extracellular recording with localized intracellular access, extending the functionality of the neuromimetic interface beyond conventional extracellular electrophysiology.

## Conclusion

The ability to engineer bioelectronic interfaces that structurally integrate with living neural tissue remains a central challenge in neurotechnology. Here, we demonstrated that the morphology of conducting-polymer electrodes can be tailored to generate dendrite-inspired architectures that regulate neuron-electrode interactions across multiple length scales. By controlling the electrodeposition process, we established design principles linking fabrication parameters to hierarchical electrode morphology and electrochemical performance, enabling the reproducible fabrication of interfaces spanning planar coatings to highly branched dendritic structures.

The engineered topography fundamentally modulated the physical interaction between neurons and the electrode surface. Whereas planar electrodes exhibited a pronounced membrane-electrode cleft, the dendritic architectures promoted conformal membrane wrapping and localized membrane invaginations, consistent with an active engulfment mechanism previously reported for engineered micro- and nanostructured interfaces. Importantly, these ultrastructural changes remained spatially confined to the neuron-electrode interface without inducing structural changes at the network level, such as altered neuronal spatial clustering or astrocyte coverage. They were accompanied by possible accumulation of the postsynaptic scaffold protein, namely PSD95, suggesting that hierarchical dendritic topography might promote synaptic organization at sites of intimate bioelectronic coupling.

Beyond structural integration, the dendrite-inspired electrodes also modulated neuronal function. SF electrodes consistently exhibited increased firing and burst rates, indicating that the enhanced membrane conformability translates into stronger functional coupling rather than simply affecting passive electrical recording. The comparatively smaller effect observed for LF electrodes is likely explained by coupling to distal positions near the tip of the branch where both branch dimensions and nanoscale roughness progressively decrease. Those observations suggest that the structural features governing membrane conformability also determine the efficiency of electrical coupling between neurons and the bioelectronic interface. Finally, we show that the same dendritic architectures support localized laser-assisted optoporation, enabling transient intracellular electrophysiological access while preserving extracellular recording capability. The ability to combine extracellular recording, localized intracellular interrogation and neuromimetic structural integration within a single organic bioelectronic platform extends the functionality of conventional neural interfaces and highlights the versatility of the dendritic electrode architecture.

Although further optimization of the fiber geometry and electrochemical performance will be required to maximize signal fidelity and intracellular signal quality, our results establish hierarchical dendritic bioelectronics as a general strategy for engineering neural interfaces that actively direct structural and functional integration with living neurons. More broadly, these findings suggest that programming the morphology of organic electronic materials provides a route towards bioelectronic systems in which artificial and biological neural networks are coupled not only through electrical communication, but also through the structural organization of the neuron-electrode interface.

## Materials & Methods

### Multi electrode array fabrication and dendritic fibers deposition

The MEAs were fabricated as previously reported^30^, in brief, using a quartz substrate with a size of 240 × 240 mm^2^ to which on top a metal stack of Ti/Au/Ti with layer thicknesses of 20/200/10 nm was deposited to pattern the electrode pads and feedlines. The electrode pads were patterned to have a size of 200 × 200 µm^2^ or 26 × 26 µm^2^. Alternating layers of 200 nm silicon oxide and 100 nm silicon nitride up to a thickness of 800 nm were deposited. Another layer of tantalum pentoxide or titanium oxide with a thickness of 40 nm was then deposited on top. Electrode openings with a diameter of 50, 24, 12 and 8 µm, were etched at the center of the electrode’s pad.

The MEAs were cleaned by immersion in a beaker filled with acetone (Th. Geyer, cat. No. 2659). The beaker was then placed for 10 minutes in an ultrasonic bath (Emmi-40 HC, Emmi EMAG AG, Germany) at a power of 50%. Afterwards, the acetone was replaced by isopropanol (Th. Geyer, 45629.02) and left in the ultrasonic bath for 10 more minutes at a power of 50%. Afterwards the sample was blow dried using a nitrogen gun. For optoporation experiments, the MEA was bonded using reflow soldering to a custom-made printed circuit board (PCB) to later allow the measurement with the MEA2100-mini head stage (Multi Channel Systems, Germany).

A glass ring with an outer diameter of 17 mm and a height of 12 mm was glued onto the center of the cleaned MEA using a biphasic glue eco-sil (picodent, cat. No. 1300 6100) by mixing the two components in a 1:1 ratio. The biphasic glue was applied to the rim of the glass ring. The glass ring was then placed on top of the MEA and left to dry at room temperature for at least 30 minutes.

To deposit the conductive dendritic fibers, a monomer solution was prepared by adding 50 mM EDOT (Sigma Aldrich, cat. No. 483028-10G) and 1 mM tetrabutylammonium hexafluorophosphate (TBAPF_6_) (Sigma Aldrich, cat. No. 281026-100G) in acetonitrile (Sigma Aldrich, cat. No. 34851-2.5L). The Arkeo measurement system (Cicci Research, Italy), consisting of two source-meter units (SMUs), was used to apply the deposition signal. The MEA was connected in a two-electrode setup, where the MEA electrode is connected as the working electrode, and a Pt-wire immersed in the monomer solution acts as the counter electrode. The Pt-wire was coiled to ensure a more homogenous electrical field. Symmetric, biphasic voltage pulses with an amplitude ranging from 3.5 to 5.5 V were applied as the deposition voltage. The signal frequency was set to between 50 and 200 Hz, and the deposition time was varied from 2 to 10 seconds. Long and short fiber electrodes on the 24 µm diameter electrodes were fabricated using a starting voltage of 2.3 V with a gradual increase of 8 mV per period until the voltage of 3.5 V was reached. The short and long fiber electrodes were deposited for a total duration of 2 seconds and 6 seconds, respectively. For the deposition of the 2D electrodes a constant current of 69.1 nA or 300 nA was applied for the 24 and 50 µm diameter electrodes, respectively.

### Electrochemical impedance spectroscopy (EIS)

The potentiostat VSP-300 (Biologic, Germany) was used to perform EIS. The investigated frequency range was set between 3 MHz and 100 Hz. To ensure the system behaves linearly, a perturbation signal with an amplitude of 10 mV was used. As a reference electrode, a pellet Ag/AgCl (Science Products GmbH, cat. No. E-206) electrode was used, and Dulbecco’s Balanced Salt Solution (DPBS) without Ca^2+^ and Mg^2+^ (Thermo Fisher, cat. No. 14190169) was used as electrolyte.

### Cyclic voltammetry (CV)

The potentiostat VSP-300 (Biologic, France) was used to perform CV measurements. The potential window was set between -0.6 and 0.6 V with a scan rate of 100 mV s^-1^. The voltage window was chosen to be below the oxidation potential of PEDOT:PF_6_. A leakless Ag/AgCl electrode (eDAQ, cat. No. ET072-1) was used as reference electrode and DPBS without Ca^2+^ and Mg^2+^ (Thermo Fisher, cat. No. 14190169).

### Culturing of primary cortical neurons

Primary cortical neurons were isolated from embryonic day 18-19 Wistar rat embryos (Javier, France). Use of primary tissues in this work has been approved by the state animal ethics committee, the Landesumweltamt für Natur, Umwelt und Verbraucherschutz Nordrhein-Westfalen, Recklinghausen, Germany, under permit number 81-02.04.2023.A172. It has been conducted according to local animal protection regulations and is reported according to the ARRIVE guidelines.

Substrates were prepared for cell culture by sterilizing and coating the surface. Ethanol absolute (99.9%; Th. Geyer, cat. No. 2246-2.5L) was mixed with DI-water to obtain a 70% v/v solution. The samples were then submerged in 70% ethanol for 15 minutes. Afterwards, the samples were submerged in DI-water for 15 minutes. The samples were coated for one hour using a solution containing Poly-L-Lysine (PLL) (Sigma Aldrich, cat. No. P1399) and Hanks′ Balanced Salt solution (HBSS; Sigma Aldrich, cat. No. H6648) in a ratio of 1:100 PLL:HBSS. The solution was added to the sample and was left for one hour. Afterwards, the samples were washed with clear HBSS 3 times.

The extracted rat cortices were directly prepared or stored for up to 1.5 weeks in Hibernate A (Thermo Scientific, cat. No. A1247501) until plating. Neuronal cells were isolated using trypsin digestion. First, the tissue was put in 2 mL of 0.05% cold trypsin EDTA (Invitrogen, cat. No. 25300-062) and then incubated at 37°C for 10 minutes. Then the tissue was washed by removing the supernatant and adding fresh Neurobasal medium (Invitrogen, cat. No. 21103-049) containing 1% B-27 (Invitrogen, cat. No. 17504-044), 0.25% L-glutamine (Invitrogen, cat. No. 25030-024) and 0.1% gentamicin (Sigma Aldrich, cat. No. G1397) three times. The tissue was then triturated until it was separated. Afterwards, the supernatant was transferred into a tube of fresh media, followed by the cell counting and plating of the desired cell number. Finally, the plated samples were transferred into an incubator at 5% CO_2_ at 37°C^31^. For extracellular electrophysiological recordings, optoporation experiments and live/dead assays, a density of 130,000 cells cm^-2^ was plated, while for immunostaining, a density of 39,000 cells cm^-2^ was plated.

2 to 4 hours after plating, the full media was exchanged, afterwards half of the media was changed every 3 to 4 days. For this, new media was prepared with Neurobasal Plus medium (Thermo Fisher, cat. No. A3653401) as base. Then adding 1% B-27 Plus Supplement (Thermo Fisher, cat. No. A3653401), 0.25% glutamax (Thermo Fisher, cat. No. 35050061) and 0.1% gentamicin (Sigma Aldrich, cat. No. G1397). Afterwards, the medium was heated in the water bath at 37°C and filtered with a 0.22 µm sterile filter (Merck, cat. No. SLGP033RS).

To reuse the MEAs after an experiment, they were cleaned by exchanging the cell media and replacing it with 0.05% Trypsin-EDTA (Invitrogen, cat. No. 25300-062). The samples were then placed on a hotplate at 37°C for 10 minutes. This step was repeated twice, followed by rinsing with 3 times deionized water. Afterwards, the samples were stored in DPBS until sterilization and coating of the samples.

### Culturing of human induced pluripotent stem cell-derived cardiomyocytes (hiPSC-CMs)

HiPSC-CMs (iCell Cardiomyocytes, cat. No. R1105) were purchased from FUJIFILM Cellular Dynamics, Inc. (FCDI). Prior to cell seeding, the devices were sterilized in 70% ethanol, prepared by mixing ethanol absolute (Sigma Aldrich, cat. No. 32221-2.5L-M) with DI-water, for 30 minutes and subsequently rinsed three times with sterile water. To prepare the surface, the electrode area was coated sequentially with 0.02% gelatin (Sigma Aldrich, cat. No. G1890) for 1 hour and 50 μg mL^-1^ fibronectin (Roche, cat. No. 11051407001) for 1 hour, both at 37°C, 5% CO_2_ in a humidified incubator. After removing the fibronectin solution, hiPSC-CMs were directly seeded onto the electrode area at a density of 30,000 cells/device in plating medium (Fujifilm Cellular Dynamics, cat. No. M1001). Two days post-seeding, the plating medium was replaced with maintenance medium (Fujifilm Cellular Dynamics, cat. No. M1003). Subsequently, 50% of the culture medium was changed with fresh maintenance medium every 2 to 3 days. Electrophysiological measurements were conducted between days 10 and 14 after seeding, following the manufacturer’s guidelines.

### Critical point drying (CPD)

Samples were taken from the incubator and then washed with DPBS without Ca^2+^ and Mg^2+^ (Thermo Fisher, cat. No. 14190169) and then washed in 0.1 M sodium cacodylate buffer, which was diluted with MiliQ water from a 0.4 M sodium cacodylate buffer (0.4 M; Science Services, cat. No. 11654) at a physiological pH for 5 minutes. The cells were fixed using a fixation buffer containing 2.5% glutaraldehyde (25%; Sigma Aldrich, cat. No. 354400) in 0.1 M sodium cacodylate for 2 hours at room temperature. Afterwards, the specimens were kept at 4°C. After a thorough wash with deionized water, samples were dehydrated in a gradient series of cold ethanol absolute (30%, 50%, 2 × 70%, 3 × 95%, and 3 × 100% in deionized water). Each step was carried out for 10 minutes at 4°C. Next, the CPD was performed using the CPD machine 931 (Tousimis, USA). The samples were loaded into the CPD chamber, and the ethanol was slowly exchanged with liquid CO_2_. After 10 to 15 exchanges, the chamber was heated up until the critical point of CO_2_ was reached. The chamber was then vented. Afterwards, the samples were sputtered with a thin layer of Pt (15 mA, 90 s).

### Scanning electron microscopy (SEM)

CPD processed samples were imaged at DIV5 using a LEO 1550 VP (Zeiss, Germany) with an acceleration voltage of 3 kV and working distance of 4.1-4.2 mm.

For the imaging of samples without cells, they were dried and then sputtered with a thin layer of iridium (15 mA, 60 s) and then imaged using the GeminiSEM 360 (Zeiss, Germany) with an acceleration voltage of 3 kV and a working distance of 7.7 - 7.8 mm.

### Ultra-thin plasticization (UTP) on neurons

UTP was performed as previously described^20^. MEAs were fixed by incubation in 2.5% glutaraldehyde in 0.1 M sodium cacodylate buffer for 2 hours, prepared as described for the CPD procedure. Samples were then washed 3 times in 0.1 M sodium cacodylate buffer (10 min per wash) at room temperature and stored in 0.1% glutaraldehyde and 0.1 M sodium cacodylate buffer. Next, the samples were quenched with freshly prepared 20 mM glycine solution in 0.1 M sodium cacodylate (20 minutes, 4°C). Afterwards, samples were rinsed 3 times at room temperature with 0.1 M sodium cacodylate, followed by post fixation in a solution containing 2% aqueous osmium tetroxide (Science Services, cat. No. 19170) and 1% potassium ferrocyanide (Electron Microscopy Sciences, cat. No. 25102-20) in 0.1 M sodium cacodylate for 1 hour at 4°C, protected from light. Subsequently, the samples were washed 3 times with ice-cold 0.1 M sodium cacodylate (3 times, 5 minutes each wash) and immersed in a filtered 1% thiocarbohydrazide (TCH; Electron Microscopy Sciences, cat. No. 21900) solution in deionized water for 20 minutes at room temperature, followed by 3 times of washing with ice-cold deionized water (5 minutes each wash). This step is followed by incubation in a 1% aqueous osmium tetroxide solution (Science Services, cat. No. 19172) in 0.1 M sodium cacodylate at room temperature for 1 hour in a dark environment. After washing with distilled water at room temperature (3 times, 5 minutes each wash), the samples were incubated overnight at 4°C in a 1% uranyl acetate (Electron Microscopy Sciences, cat. No. 22400-2) solution in deionized water. Following washing with ice-cold distilled water (5 times, 5 minutes each wash), the specimens were then incubated in a 0.15% tannic acid solution (Electron Microscopy Sciences, cat. No. 21710) in deionized water for 3 minutes at 4°C. After a thorough wash with deionized water, samples were dehydrated in a gradient series of cold ethanol absolute (30%, 50%, 2 × 70%, 3 × 95%, and 3 × 100% in deionized water). Each step was carried out for 10 minutes at 4°C. For resin embedding, the samples were gradually infiltrated with low-viscosity embedding media Spurr’s kit (Electron Microscopy Science, cat. No. 1430) using increasing ethanol to resin ratios. The initial embedding used a 3:1 ethanol/deionized water to resin ratio for 2 hours at room temperature, followed by a 2:1 ethanol-to-resin ratio for 2 hours at room temperature, and a 1:1 ethanol-to-resin ratio, overnight at room temperature. The following day, embedding was performed with a 2:1 and 3:1 resin-to-ethanol ratio for 2 hours each embedding step, followed by pure fresh resin incubation overnight and throughout the following day (pure resin, 2 times with each step lasting 2 hours). Samples were then placed in a vertical position for 3 hours to remove the excess resin. Resin was polymerized for 24-70 hours, at 70 °C. Afterwards, samples were mounted on aluminum stubs using silver conductive paste.

### Focused ion beam – Scanning electron microscopy (FIB-SEM) on neurons

The cross-section UTP processed samples were obtained using the SEM FEI Helios Nanolab 600 (Thermo Fisher Scientific, United States). Prior to imaging, specimens were sputtered with a layer of iridium (15 mA, 60 s). The specimens were loaded into the dual-beam vacuum chamber (Thermo Fisher, Helios CX5). The region of interest (ROI) was identified, and a layer of platinum (1 - 2 µm thickness, current 1.4-22 nA, voltage 2-3 kV) was applied *via* ion beam deposition. Afterward, the stage was tilted to 52° and a layer of platinum (1 - 1.5 µm thickness, current 2.5-9.3 nA, voltage 30 kV) was applied *via* ion beam deposition and bulk milling was carried out with the same settings, with a thickness of 8-15 µm, depending on the thickness of the resin. Cross-sections were created by trenching with the ion beam (7-10 µm, current: 9.3 nA, voltage: 30 kV), followed by polishing the interface with the ion beam (current: 0.08 - 0.43 nA, voltage: 30 kV). Finally, scanning electron micrographs were acquired in backscattered mode with a through-lens detector (TLD) at high resolution with electron beam set to 3 kV and 0.08-0.43 nA with a working distance of 4.1 mm.

### Immunohistochemistry on neurons

Primary cortical neurons were fixed on DIV10 and DIV22 using a buffer based on deionized water with the addition of 80 mM PIPES (Merck, cat. No. P7643), 5 mM ethylene glycol-bis[β-aminoethyl ether]-N,N,N’,N’-tetraacetic acid (EGTA) (Sigma Aldrich, cat. No. E3889), 2 mM MgCl_2_ (Roth, cat. No. A537.1). The pH was set to 6.8. Afterwards the solution was filtered with a 0.22 µm sterile filter (Merck, cat. No. SLGP033RS). A 16% paraformaldehyde (PFA) (Thermo Fisher, cat. No. 1150058) stock solution was diluted with the buffer to achieve a 4% PFA buffer. The samples were then taken from the incubator, and the medium was replaced with the 4% PFA buffer. The samples were then left for 6 to 7 minutes to preserve synaptic targets. Afterwards, the samples were washed with DPBS without Mg^2+^ and Ca^2+^ (Thermo Fisher, cat. No. 14190169) 3 times for 5 minutes each. For immunostaining, the neurons were permeabilized and blocked for 1 hour in a buffer containing 0.2% Triton X-100 (Sigma Aldrich, cat. No. X100) and 10% goat serum (Fisher Scientific, cat. No. 16-210-072) in DPBS without Ca^2+^ and Mg^2+^. Primary and secondary antibodies were diluted in 5% goat serum, 0.1% Triton X-100 in DPBS. The primary antibodies goat anti-mouse postsynaptic density protein-95 (PSD95, 1:250; Biozol, cat. No. ANI-75-028), goat anti-rabbit microtubule-associated protein 2 (MAP2, 1:250; Merck, cat. No. AB5622) and goat anti-chicken glial fibrillary acidic protein (GFAP, 1:250; Abcam, cat. No. ab134436) were incubated at 10°C overnight. Afterwards, the samples were washed with DPBS without Mg^2+^ and Ca^2+^ (3 times, 5 minutes each). Next, the specimens were incubated with the secondary antibodies goat anti-mouse IgG(H+L) highly cross adsorbed secondary antibody alexa fluor plus 488 (1:500; Thermo Scientific, cat. No. A32723), goat anti-rabbit IgG(H+L) secondary antibody alexa fluor 568 (1:500; Thermo Scientific, cat. No. A11036) and goat anti-chicken IgY(H+L) cross adsorbed secondary antibody alexa fluor plus 647 (1:500; Thermo Scientific, cat. No. A32933). Afterwards, they were washed with DPBS Mg^2+^ and Ca^2+^ (3 times, 5 minutes each). Finally, the samples were mounted using a mounting medium including DAPI (Bitoium, cat. No. 23002), placing a 13 mm coverslip (Fisher Scientific, cat. No. 10513234) on top of the area with the cells and sealing with nail polish. The mounted samples were then stored at 4°C until imaging.

Widefield fluorescence microscopy was performed on an AxioImager.M2 (Zeiss, Germany) equipped with a Viluma LED illumination system and a AxioCam 820. Images and image stacks were acquired in the ZEN 3.12 software (Zeiss, Germany). The following reflectors were used: DAPI (420 - 470 nm), eGFP (500 - 550 nm), DsRed (570 - 640 nm), Cy5 (662-700 nm), and DIC reflected light. Where indicated, an Apotome 3 grid (Zeiss, Germany) was inserted into the optical path. During this acquisition, the grid is tilted back and forth in the light path and projected onto the specimen. The different phase images are then combined to reduce the out-of-focus light.

Optical micrographs of the whole-electrode arrays were acquired with a 20×/0.8 air objective using the tiling function with a 10% overlap to acquire a composite area of 60 × 60 mm² centered on the electrode region. Fluorescence micrographs were acquired using the following excitation and acquisition settings: DAPI (385 nm, 15% intensity, 70 ms exposure), DsRed (567 nm, 25% intensity, 70 ms exposure), Cy5 (630 nm, 8% LED intensity, 70 ms exposure) and reflected-light brightfield (385 nm, 430 nm, 475 nm, 567 nm, 630 nm and 735 nm, intensity of 0.2% each, 2 ms exposure) with a bit depth of 14 bit and a pixel size of 0.137 µm.

Optical Micrographs of single electrodes were acquired using a 40×/1.4 oil objective with the immersion oil 518 F (Zeiss, cat. No. 444960-0000-000) as transmission medium between objective and specimen. For each position the DAPI (385 nm, 15% LED intensity, 200 ms exposure) and reflected-light brightfield (385 nm, 430 nm, 475 nm, 567 nm, 630 nm and 735 nm, intensity of 0.2% each, 20 ms exposure) channel were acquired in the focus plane, set by the software autofocus in reference to the reflected light brightfield channel. Additionally, the eGFP (475 nm, 40% intensity, 150 ms exposure), DsRed (567 nm, 25% intensity, 70 ms exposure) and Cy5 (630 nm, 8% intensity, 70 ms exposure) were acquired using the Apotome 3 function (5 phase images) and Z-stack of 5 µm (0.275 µm step) centered around the focus plane of the DsRed channel, which was set by using the software autofocus. Both acquisitions have a bit depth of 14 bits, and a pixel size of 0.068µm.

### Biocompatibility assay

Biocompatibility assays were performed on primary cortical neurons at DIV2. The staining solution was prepared by adding 10% Ethidium homodimer (Sigma Aldrich, cat. No. E1903) and 10% Calcein AM (Invitrogen, cat. No. C3099) to Neurobasal medium (Invitrogen, cat. No. 21103-049). 10 µL of the staining solution was added to the samples and mixed by pipetting up and down. The samples were then left to incubate for 10 minutes. Afterwards, the samples were imaged using the fluorescence microscope Axio Imager.Z1 (Zeiss, Germany). For illumination, an HXP-120 halogen lamp with the following reflectors was used: GFP (500-550 nm) and DsRed (570-640 nm). For the acquisition, a 10×/0.3 water immersion objective was used. In each position, fluorescence micrographs were acquired from the GFP (494 nm, 50 ms) and DsRed (550 nm, 100 ms exposure). All images have a bit depth of 12 and a pixel size of 0.645 µm. Micrographs were acquired up to a maximum of 30 minutes after acquisition.

### Immunocytochemistry on cardiomyocytes

On day 14 after seeding hiPSC-CMs, immunofluorescence labeling for cardiac troponin T (cTnT) was performed using the Human Cardiomyocyte Immunocytochemistry Kit (Life Technologies, cat. No. A25973). Cells were initially fixed with 4% (w/v) formaldehyde in Dulbecco’s Phosphate Buffered Saline (DPBS) (Life Technologies, cat. No. A24344) for 15 minutes at room temperature, permeabilized with 1% saponin (Life Technologies, cat. No. A24878) for 15 minutes and blocked with 3% bovine serum albumin (BSA) (Life Technologies, cat. No. A24353) for 30 minutes. Samples were then incubated with a mouse anti-cTnT/TNNT2 primary antibody (Life Technologies, cat. No. A25969; 1:1000) overnight at 4°C. Following three washing steps in washing buffer (Life Technologies, cat. No. A24348), the cells were incubated for 1 hour at room temperature with a donkey anti-mouse Alexa Fluor 488 secondary antibody (Life Technologies, cat. No. A25972; 1:250). Finally, nuclei were counterstained with DAPI (Life Technologies, cat. No. R37606). Micrographs were acquired using an upright fluorescence microscope (Eclipse FN1, Nikon) equipped with a 16-bit Hamamatsu ORCA-Flash4.0 sCMOS camera (C13440-20CU, Hamamatsu Photonics) at full frame resolution (2048×2048 pixels, binning 1×1). Low-magnification overview micrographs were captured using a 20×/0.4 air objective (working distance 20 mm), while high-magnification cellular details were acquired using a 60×/1.0 water immersion objective (working distance 2.8 mm). These acquisitions were performed with a bit depth of 16 bit and spatial resolutions (pixel sizes) of 0.325 µm/pixel for the 20× objective and 0.1083 µm/pixel for the 60× objective, respectively. For each field of view, fluorescence signals were detected using standard filter sets for Alexa Fluor 488 channel (Ex/Em 495/519 nm) and DAPI channel (Ex/Em 358/461 nm). Transmitted light brightfield micrographs were also acquired to correlate cellular immunostaining with the electrode array. Exposure times and illumination intensities were optimized per channel to maximize signal-to-noise ratio while avoiding pixel saturation (1.0 - 2.0 ms for brightfield and 1.0 - 4.0 s for fluorescence channels) and were kept strictly constant across all acquired samples. Post-acquisition processing was performed using ImageJ/Fiji software. Where necessary to reduce non-uniform background fluorescence and improve the signal-to-noise ratio, a Fast Fourier Transform (FFT) Bandpass Filter was applied (filtering spatial structures larger than 200 pixels) while preserving original pixel intensity dynamics. Otherwise, linear brightness and contrast adjustments were performed manually to optimize visual clarity while strictly avoiding signal saturation.

### Ultra-thin plasticization (UTP) on cardiomyocytes

The samples were prepared following the ultra-thin plasticization procedure previously described^32^. Biological samples were fixed in 2% (v/v) glutaraldehyde (Società italiana chimici, cat No. 16220) prepared in 0.1 M sodium cacodylate (Società italiana chimici, cat. No. 11655) buffer (pH 7.4) by incubation either overnight at 4°C or for 1 hour at room temperature. Following fixation, the specimens were washed three times for 10 min each in 0.1 M sodium cacodylate buffer before proceeding with the subsequent processing steps.

The buffer was then replaced with 20 mM glycine (Sigma-Aldrich, cat. No. T6522-100MG) prepared in 0.1 M sodium cacodylate buffer, and the samples were incubated for 20 minutes. The specimens were subsequently incubated in a solution containing 4% (v/v) aqueous osmium tetroxide (Società italiana chimici, cat. No. 19190) and 2% potassium ferrocyanide (Electron Microscopy Sciences, cat No. 25102-20) for 1 hour at 4°C, protected from light. After three washes with 0.1 M sodium cacodylate buffer, a second incubation was carried out in 2% (v/v) aqueous osmium tetroxide (Società italiana chimici, cat. No. 19190) for 30 minutes at room temperature. The samples were rinsed with deionized (DI) water and immersed in 1% filtered thiocarbohydrazide (TCH; Sigma-Aldrich, cat. No. 223220-5G) prepared in DI water for 20 minutes at room temperature, followed by three rinses with DI water. They were then incubated overnight at 4°C in 4% (v/v) aqueous uranyl acetate (SIC, cat. No. 22400-4) for en bloc staining. After three additional rinses with DI water, the samples were incubated in 0.15% (v/v) tannic acid for 3 minutes at 4 °C. Dehydration was performed through a graded ethanol series (30%, 50%, 75%, 2× 95%, and 100% ethanol), with each step lasting 10 minutes at 4°C. Absolute ethanol was then replaced twice at room temperature. The samples were gradually embedded in resin consisting of 25 mL NSA, 8 mL D.E.R. 736, 10 mL ERL 4221, and 301 μL DMAE (Società italiana chimici, cat. No. 14300) using increasing resin concentrations. The specimens were first embedded in a 3:1 ethanol-to-resin mixture for 2 hours, followed by a 2:1 mixture for a further 2 hours, and finally in pure resin overnight and throughout the following day. The samples were then positioned vertically for 2 hours and polymerized at 70°C for 24 hours. For electron microscopy imaging, the specimens were mounted onto 3.2 mm diameter aluminum stubs using silver conductive paste (RS Pro, cat. No. 123-9911) and coated with a 15 nm layer of gold prior to imaging.

### Focused ion beam – Scanning electron microscopy (FIB-SEM) on cardiomyocytes

The specimens were introduced into the vacuum chamber of a dual-beam FIB-SEM instrument (Thermo Fisher Helios CX 5), and the region of interest (ROI) was identified. A protective platinum layer (0.5 µm thick) was deposited by ion beam-induced deposition using a 30 kV accelerating voltage and a beam current of 0.43 nA. Cross-sections were then prepared by trench milling with the ion beam (4 µm cutting depth, 30 kV, 0.79 nA), followed by a polishing step performed at 30 kV using beam currents between 0.23 and 0.43 nA. Scanning electron micrographs were acquired in backscattered electron mode at 3 kV and 0.17 nA, using a dwell time of 10 µs and high-resolution imaging settings.

### Electrophysiological recordings and optoporations

For extracellular electrophysiological recordings on primary neurons, the biomas system (FZJ, Germany) was used. It allows for simultaneous recordings of 64 electrodes. The MEA can be connected to the system via a custom designed head stage. Electrophysiological recordings were performed on DIV10, DIV14 and DIV22. An Ag/AgCl electrode was placed into the cell media to provide a stable reference potential for the measurement. Recordings were performed with a sampling rate of 10,000, a voltage gain of 1,000 and a high pass filter with a cutoff frequency of 0.1 Hz. To obtain extracellular electropysiological recordings in combination with optoporations on neurons, the Intracell system (Foresee Biosystems, Italy) was used on primary cortical neurons on DIV14. To porate the membrane a train of fast laser pulses with a wavelength of 976 nm and average laser power of 14 mW was applied. The laser is pulsed with a frequency of 1 MHz, and the total pulse train duration was set to 10 milliseconds. Simultaneously, the extracellular activity was recorded using a ME2100 system with the MEA2100-mini head stage (Multi Channel Systems, Germany). The sampling rate was set to 25,000, and a hardware high pass filter of 0.1 Hz and low pass filter of 3.5 kHz was used.

Extracellular electrophysiological recordings on hiPSC-CMs were carried out at 37°C outside the incubator. The culture medium was replaced with fresh maintenance medium at least 2 hours prior to measurements. The extracellular field potentials and intracellular action potentials were acquired using a MEA2100-Mini-System (Multi Channel Systems, Germany). Optoporation was performed according to the methodology described previously^33^. A solid-state Nd:YAG laser (Plecter Duo, Coherent) operating at its fundamental wavelength (λ = 1064 nm), with an 8 ps pulse width and an 80 MHz repetition rate, served as the excitation source. The laser beam was directed through a custom-modified upright microscope (Eclipse FN1, Nikon) integrated with the MEA2100-Mini-System. A 20× air objective (numerical aperture 0.4, working distance 20 mm) was used to observe the cardiomyocytes on the array and to focus the NIR light for membrane poration. The average laser power after the objective was approximately 1 mW.

### Atomic force microscopy (AFM)

The measurements were performed using a Dimension Icon AFM (Bruker Corporation, USA) equipped with ScanAsyst-Fluid cantilevers (Bruker, USA). Imaging was carried out in air under ambient conditions using ScanAsyst mode. The scanned area was adjusted depending on the substrate and sample morphology. For gold, 2D electrodes, and SF, measurements were performed in tapping mode with a scan rate of 1 Hz. For LF, tapping mode was operated at a scan rate of 2 Hz.

## Image analysis

### Biocompatibility assay

Raw image files were batch-processed using a custom macro in ImageJ/Fiji^34^ (NIH, USA). The raw images were first separated into separate channels and then converted into a 16 bit format. Afterwards, the brightness and contrast were adjusted uniformly using a predefined look up table. Segmentation of live and dead cells was performed by global thresholding with fixed intensity ranges (live: 125-255, dead: 100**-**225). The threshold selection was empirically determined and maintained constant throughout the analysis. Binary masks were generated for each channel and subjected to watershed segmentation to separate touching or overlapping cells. Afterwards, objects were filtered based on morphological criteria. For live cells, size and circularity ranges were set from 0**-**200 pixels and 0.25**-**1, respectively. For dead cells, ranges were set to 2**-**100 pixels and 0.5**-**1. Quantification was performed using the built-in particle analysis tool in ImageJ/Fiji. The number of live and dead cells was determined for each field of view. Finally, the live fraction was determined by the ratio of live cells over the total number of cells. All analysis parameters were held constant across samples.

For display, the brightness and contrast of were adjusted in ImageJ/Fiji setting ranges for EtHD (100-3500) and CalceinAM (100-3500).

### Immunohistochemistry on neurons

Images that were acquired in tiles were stitched using the ZEN 3.12 software (Zeiss, Germany) and the MAP2 (DsRed) channel as reference for the stitching. For the single electrode acquisitions, first the Apotome processing was performed using the Zen pro (3.12) software (Zeiss, Germany), afterwards the images were converted into maximum intensity projections. Finally, the acquisitions of the DAPI and reflected-light brightfield channels were merged with the acquisitions of the eGFP, DsRed and Cy5 channels.

To enable fast and uniform processing, every image was converted to 8-bit using a contrast-stretching step. The analysis was performed on two complementary datasets. The first comprised whole-array mosaic images; then a subset of tiles with clearly identifiable electrodes was selected for analysis. The second dataset consisted of single-electrode scenes, acquired at higher magnification, and was used specifically for PSD-95 puncta detection and orientation analysis. To convert these images into quantifiable biological data, a set of custom image processing routines was used, that draw boundaries around structures of interest and label them as distinct objects. The same core segmentation logic was applied across both datasets to ensure consistency.

For the synaptic marker PSD-95, synaptic proteins were identified as roughly circular bright puncta using blob detection (OpenCV’s SimpleBlobDetector). To enhance signal contrast, a bilateral filter was first applied to suppress camera noise without blurring sharp edges, followed by a morphological top-hat transformation with a large structuring element (radius 20 pixels) to flatten the background illumination. Detection was then performed on the background-subtracted image using a step-wise multi-threshold strategy (starting at a low intensity threshold of 7, with a step of 20). Candidate puncta were constrained using physical size filters (equivalent diameter between 0.30 µm and 1.20 µm; maximum area 1.2 µm²), as well as shape filters including circularity (≥ 0.3), which preserved compact nascent puncta, and inertia ratio (≥ 0.1), a measure of elongation that allowed retention of larger, more complex puncta characteristic of mature synapses. This combination of size and morphological constraints aims to exclude non-specific debris and capture the morphological diversity of PSD-95 puncta across developmental timepoints. This blob-based approach was chosen for its high computational speed when processing large images.^35^

For dendritic (stained with MAP2) and astrocytic processes (stained with GFAP), we used the same segmentation pipeline for both markers. This segmentation pipeline first applied a local contrast enhancement algorithm called CLAHE (Contrast Limited Adaptive Histogram Equalization) to compensate for uneven staining or illumination across the image. After reducing background haze with a second top-hat filter, the algorithm used an adaptive thresholding step, meaning the brightness cutoff used to separate signal from background was calculated dynamically for each small neighborhood of pixels, rather than applying a single global value. The resulting binary outlines were then processed through morphological closing, a routine that bridges small gaps, producing continuous representations of neuronal and glial processes.^36,37^

Nuclei, visualized with DAPI, were segmented using a classical watershed algorithm preceded by an Otsu threshold. Otsu’s method finds the optimal global intensity threshold to separate bright nuclei from the darker background. The watershed step then separates nuclei that are touching or overlapping.

Electrodes were localized manually within the brightfield channel. Initially, Otsu thresholding was applied to the brightfield image to create a binary map of dark (electrode) versus bright (insulating) regions; an operator then visually inspected the map and manually selected the connected component that corresponded to each electrode, yielding precise binary masks for the electrode positions.

Following segmentation, binary masks and object feature tables were generated. For the images of the whole electrode-array, the MAP2, GFAP, and DAPI masks were overlaid to classify each nucleus as neuronal (MAP2 positive) or astrocytic (GFAP positive). Therefore, for each detected nucleus, the fraction of its area overlapping with MAP2 and the fraction overlapping with GFAP were computed^38,39^. A nucleus was classified as a neuron if the MAP2 overlap fraction was at least 0.3 and the GFAP overlap fraction was below 0.3; as an astrocyte if the GFAP overlap fraction was at least 0.3 and the MAP2 fraction was below 0.3; and as “other” otherwise. From these classified nuclei, those located within an 80 µm radius of the electrode center were used to compute the probability of finding a neuron within the radius , the number of neurons divided by the sum of neurons and astrocytes within that radius, and the astrocyte fraction within the radius, the number of astrocytes divided by the same sum. Both metrics are unit-less proportions ranging from 0 to 1, reflecting the local cellular composition immediately surrounding the electrode.

For the single-electrode images, PSD-95 puncta that overlapped simultaneously with the MAP2 mask and the electrode mask were counted. This raw count was normalized to the electrode mask area (in µm²) to obtain the electrode-specific PSD-95 density, expressed as puncta per µm² of electrode surface. Bar plots show the mean electrode-normalized density ± SEM per condition and DIV, providing a standardized measure of synaptic puncta localized specifically to the electrode-cell interface.

Additionally, dendrite masks were skeletonized and decomposed into individual branch segments^40,41^. For each segment, the angular bearing relative to the electrode center was measured, mapping where the dendritic mass is positioned around the electrode. From these length-weighted bearings, the alignment coefficient R was derived as the length of the mean resultant vector; it ranges from 0 (dendrites uniformly spread in a diffuse halo or bipolar arrangement) to 1 (mass concentrated in a single compass direction). Bar plots present the mean R ± SEM per condition and DIV.

For display, the brightness and contrast were adjusted in ImageJ/Fiji. Whole array images were processed setting ranges for the brightfield (4,000 -16,000), MAP2 (100 - 3,500), GFAP (100 - 4,000) and DAPI (50 - 500). Afterwards, the images were resized while keeping a DPI of 900, followed by the conversion to RGB. Single electrode images were processed setting ranges for the brightfield (1000 - 16,000), DAPI (150-2800), PSD95 (60-900), MAP2 (20-850) and GFAP (10-1500) channels, followed by the conversion to RGB.

### Atomic Force Microscopy (AFM)

FM data were analyzed with Gwyddion (v2.62). Prior to analysis, images were processed using a three-point plane correction procedure to correct for sample tilt. The root mean square (RMS, Rq) roughness was evaluated on selected regions of interest (ROIs). For flat surfaces (gold and 2D electrodes), three square ROIs of 2×2 µm² were randomly picked for each sample and averaged to obtain representative roughness values. The same ROI size was used to assess roughness along both short and long fibers, ensuring consistency across different sample types. To evaluate the roughness along the short-fiber and long fiber-branches a custom python script was used. To keep values consistent, the RMS roughness was extracted from 2×2 µm² squares placed along the branches. The distance between each consecutive square was calculated to extract the roughness along the length of the fiber branches.

### Morphological analysis

The morphology of fiber electrodes was analyzed using ImageJ/Fiji with the NeuronJ^42^ plugin. Structural features were classified analogously to neuronal morphology, with the electrode opening defined as the soma and extending fibers treated as neurite-like processes. Processes emerging directly from the electrode opening were classified as primary, those originating from primary processes as secondary, and subsequent extensions as tertiary. In cases where a process bifurcated into segments of comparable length, both segments were assigned to the same hierarchical level.

The projected area of fiber electrodes was quantified using ImageJ/Fiji. Images were first converted to binary format, after which the electrode outline was created using the wand tracing tool. The enclosed region was then used to calculate the total electrode area.

To estimate the soma-equivalent area, a circular region of interest was centered on the electrode opening and expanded to the starting point of the fiber branches. The measured area of this region was defined as the equivalent soma area.

## Data Analysis

### Extracellular electrophysiology

Electrophysiological recordings were analyzed using the spikeinterface (v0.104.5) library in Python (v3.11). As preprocessing, the single files were concatenated to achieve a single recording of 15 minutes. Noisy and dead channels were removed from further analysis. Afterwards, common mode referencing and a bandpass filter (300 Hz-4 kHz) were applied to the traces. The preprocessed traces were then fed into the spike sorter mountainsort4, using a threshold of 4.5, adjacency radius of 50 µm and the option to detect positive and negative spikes. The detected units were then subjected to threshold-based curation. In detail, only units with a SNR above 5, an ISI violation ratio below 0.2, a presence ratio above 0.8 and a firing rate between 0.1 Hz and 200 Hz were kept. The remaining units were then clustered according to their shape and units originating from non-biological signals were removed.

From each unit, the firing rate, noise level and SNR were extracted, as well as additional quality metrics including the ISI violation ratio, presence ratio, amplitude cutoff, d prime and isolation distance. Bursts were detected using a window of a maximum of 100 ms between spikes and a minimum number of 3 spikes per burst. Afterwards, the burst rate, the mean burst duration and the mean number of spikes per burst were calculated.

For unit clustering, waveforms were initially normalized by amplitude. Then waveform features including the peak-to-through duration, half-width, repolarization slope, recovery slope, peak-to-through amplitude ratio, waveform asymmetry, normalized through and peak amplitudes, the number of positive and negative waveform peaks, waveform spread, exponential decay constant, onset to through duration and baselines flatness were extracted. To identify groups of units, a two-dimensional embedding was generated using Uniform Manifold Approximation and Projection (UMAP). Clustering withing the embedded feature space was subsequently performed using Hierarchical Density-Based Spatial Clustering of Applications (HDBSCAN), which identifies clusters based on local point density while allowing units that do not belong to any cluster to be classified as noise.

### Electrochemical characterization

The fitting of the EIS data was performed using a custom python script based on the deareis library. To fit the data a randles circuit consisting of a solutions resistance (R_s_), double layer capacitance (C_dl_), charge transfer resistance (R_ct_) and reflective boundary Warburg impedance (W_o_) was used. Starting values were manually determined and then given to automatically fit the data. Graphs were plotted using a custom python script.

## Statistical analysis

### Cell experiments

For the extracellular electrophysiological recordings, immunohistochemistry and live/dead assay 12 MEAs in total were prepared, where 3 MEAs each were fully covered in one of the electrode types, either plain gold, 2D, short-fiber or long-fiber electrodes. Additionally, for the live/dead assay and immunohistochemistry 3 and 2 cover glasses were added per experiment as control, respectively. For each investigated DIV, at least 3 independent experiments were performed. For the optoporation experiments, MEAs with all electrode types on one MEA were prepared. For each independent experiment, 4-10 MEAs were prepared.

### Electrode characterization

For the characterization of plain gold, 2D, short fiber and long fiber electrodes through the means of EIS, CV, and the morphological analysis, 3 MEAs with 5 electrodes per electrode type on each were used.

AFM was performed on 3 separate MEAs with 3 electrodes per electrode type each.

### Electrophysiology

Statistical tests were performed using an ANOVA test, combined with a post-hoc Tukey’s test to determine the pairwise significance inside of a group.

### Immunostainings

Statistical analysis respected the hierarchical layout of the experiment. Analysis was performed directly on raw tile level measurements to preserve granularity while accounting for dependencies between tiles from the same image file (treated as correlated via a random intercept). For continuous scalar features, a Kruskal-Wallis test followed by a pairwise Mann-Whitney U test was performed.

For angular orientation data, to visualize the angles in the full 0-360° directional distributions, normalized circular histograms were arranged in a grid (DIVs × electrode types), with dendrite and electrode angles overlaid per subplot. Each channel was scaled to its own maximum bin height, enabling direct qualitative comparison of angular profiles across experimental groups while preserving the relative shape of the distribution. Additionally, the circular standard deviation was calculated. Statistics were performed using a Kruskal-Wallis test followed by a pairwise Mann-Whitney U test.

## Acknowledgements

F.S., F.D.E., and E.M. disclose support for the research of this work from the European Research Council starting grant BRAIN-ACT [grant number: 949478]. E.M. discloses the support of a scholarship from the German Academic Exchange Service (DAAD). The authors thank Marko Banzet for the fabrication of MEAs. The authors thank Bettina Breuer and Vanessa Maybeck for the preparation of the primary cortical neurons. The authors thank the Helmholtz Nano Facility (HNF) at Forschungszentrum Jülich for facilitating the microfabrication of the MEAs, SEM/FIB experiments, AFM and acquisitions with the support of Anja Zass. The authors thank Anoushka Jain for guidance on spike sorting. The authors thank Professor Harold MacGillavry for providing the anti-PSD95 antibody for preliminary experiment of PSD95 testing.

## Author contributions

K.L. performed the electrodepositions on the MEAs and the electrochemical characterization and further cultured, performed the experiments for the electrophysiological recordings, immunostainings, optoporations on neurons, CPD, analyzed and interpreted the data. E.M. analyzed the immunostaining images. F.D.E performed the EIS fittings and AFM analysis N.S. supported in cell culture and immunostaining, for the establishment and optimization of the neuronal immunostaining protocols. E.B.R. performed FIB-SEM cuts on all samples with neurons. C.F. conducted and analyzed the biocompatibility assay. C.L.B. supported SEM/FIB experiments on cardiomyocytes. S.M. supported the analysis of the extracellular electrophysiological recordings and immunostaining data. F.D.A and G.I. planned and performed preliminary optoporation experiments on cardiomyocytes. M.D. advised on the experimental parameters regarding the optoporation on neurons. V.C. supported electrodeposition experiments. F.S. conceived the idea, supervised the project, and the interpretation of the results and was responsible for fundraising. All authors have revised the manuscript.

## References

1. Szarowski, D. H. et al. Brain responses to micro-machined silicon devices. Brain Res. 983, 23–35 (2003).

2. Polikov, V. S., Tresco, P. A. & Reichert, W. M. Response of brain tissue to chronically implanted neural electrodes. J. Neurosci. Methods 148, 1–18 (2005).

3. Cogan, S. F. Neural Stimulation and Recording Electrodes. Annu. Rev. Biomed. Eng. 10, 275–309 (2008).

4. Conducting Polymers for Neural Prosthetic and Neural Interface Applications - Green - 2015 - Advanced Materials - Wiley Online Library. https://advanced.onlinelibrary.wiley.com/doi/full/10.1002/adma.201501810.

5. Ludwig, K. A., Uram, J. D., Yang, J., Martin, D. C. & Kipke, D. R. Chronic neural recordings using silicon microelectrode arrays electrochemically deposited with a poly(3,4-ethylenedioxythiophene) (PEDOT) film. J. Neural Eng. 3, 59–70 (2006).

6. Conducting Polymers for Neural Prosthetic and Neural Interface Applications - Green - 2015 - Advanced Materials - Wiley Online Library. https://advanced.onlinelibrary.wiley.com/doi/full/10.1002/adma.201501810.

7. Buzio, M. et al. 3D micropatterning of PEDOT:PSS/Gelatin conductive hydrogels via two-photon lithography for soft bioelectronics. *Npj Flex*. Electron. 10, 19 (2026).

8. Ojovan, S. M. et al. A feasibility study of multi-site,intracellular recordings from mammalian neurons by extracellular gold mushroom-shaped microelectrodes. Sci. Rep. 5, 14100 (2015).

9. Onesto, V. et al. Nano-topography Enhances Communication in Neural Cells Networks. Sci. Rep. 7, 9841 (2017).

10. Marinaro, G. et al. Networks of neuroblastoma cells on porous silicon substrates reveal a small world topology. Integr. Biol. 7, 184–197 (2015).

11. Stajković, N. et al. Bioinspired organic materials for seamless neurohybrid interfaces: from material design to living electronics. Mater. Horiz. 10.1039/d6mh00280c (2026) doi:10.1039/d6mh00280c.

12. Latte Bovio, C., et al. How Neuromorphic Microstructures Control In Vitro Early-Stage Neuronal Outgrowth. Adv. Sci. 13, e10822 (2026).

13. Latte Bovio, C., et al. How Neuromorphic Microstructures Control In Vitro Early-Stage Neuronal Outgrowth. Adv. Sci. 13, e10822 (2026).

14. Yang, X. et al. Bioinspired neuron-like electronics. Nat. Mater. 18, 510–517 (2019).

15. Janzakova, K. et al. Analog programing of conducting-polymer dendritic interconnections and control of their morphology. Nat. Commun. 12, 6898 (2021).

16. Ciccone, G. et al. Growth and design strategies of organic dendritic networks. Discov. Mater. 2, 7 (2022).

17. Influence of PEDOT:PSS Coating Thickness on the Performance of Stimulation Electrodes. https://advanced.onlinelibrary.wiley.com/doi/epdf/10.1002/admi.202000675?saml_referrer doi:10.1002/admi.202000675.

18. Cucchi, M. et al. Directed Growth of Dendritic Polymer Networks for Organic Electrochemical Transistors and Artificial Synapses. Adv. Electron. Mater. 7, 2100586 (2021).

19. PEDOT:PSS-coated platinum electrodes for neural stimulation | APL Bioengineering | AIP Publishing. https://pubs.aip.org/aip/apb/article/7/4/046117/2926407/PEDOT-PSS-coated-platinum-electrodes-for-neural.

20. Revealing the Cell–Material Interface with Nanometer Resolution by Focused Ion Beam/Scanning Electron Microscopy | ACS Nano. https://pubs.acs.org/doi/10.1021/acsnano.7b03494.

21. PSD-95 and PSD-93 Play Critical But Distinct Roles in Synaptic Scaling Up and Down | Journal of Neuroscience. https://www.jneurosci.org/content/31/18/6800.

22. Wagenaar, D. A., Pine, J. & Potter, S. M. An extremely rich repertoire of bursting patterns during the development of cortical cultures. BMC Neurosci. 7, 11 (2006).

23. Chiappalone, M., Bove, M., Vato, A., Tedesco, M. & Martinoia, S. Dissociated cortical networks show spontaneously correlated activity patterns during in vitro development. Brain Res. 1093, 41–53 (2006).

24. Buccino, A. P. et al. SpikeInterface, a unified framework for spike sorting. eLife 9, e61834 (2020).

25. Intracellular and Extracellular Recording of Spontaneous Action Potentials in Mammalian Neurons and Cardiac Cells with 3D Plasmonic Nanoelectrodes | Nano Letters. https://pubs.acs.org/doi/10.1021/acs.nanolett.7b01523.

26. Membrane Poration Mechanisms at the Cell–Nanostructure Interface - Dipalo - 2019 - Advanced Biosystems - Wiley Online Library. https://onlinelibrary.wiley.com/doi/10.1002/adbi.201900148.

27. Fendyur, A., Mazurski, N., Shappir, J. & Spira, M. E. Formation of Essential Ultrastructural Interface between Cultured Hippocampal Cells and Gold Mushroom-Shaped MEA-Toward “IN-CELL” Recordings from Vertebrate Neurons. Front. Neuroengineering 4, (2011).

28. Melikov, R. et al. Longitudinal and Noninvasive Intracellular Recordings of Spontaneous Electrophysiological Activity in Rat Primary Neurons on Planar MEA Electrodes. Adv. Mater. 37, 2412697 (2025).

29. Dipalo, M. et al. Intracellular and Extracellular Recording of Spontaneous Action Potentials in Mammalian Neurons and Cardiac Cells with 3D Plasmonic Nanoelectrodes. Nano Lett. 17, 3932–3939 (2017).

30. Abu Shihada, J., et al. Highly Customizable 3D Microelectrode Arrays for In Vitro and In Vivo Neuronal Tissue Recordings. Adv. Sci. 11, 2305944 (2024).

31. Brewer, G. J., Torricelli, J. R., Evege, E. K. & Price, P. J. Optimized survival of hippocampal neurons in B27-supplemented neurobasal^TM^, a new serum-free medium combination. J. Neurosci. Res. 35, 567–576 (1993).

32. Bovio, C. L., Mollo, V., Mariano, A. & Santoro, F. Electron Microscopy of Neurons on Biomimetic Substrates. in Neuronal Morphogenesis: Methods and Protocols (ed. Toyooka, K.) 11–20 (Springer US, New York, NY, 2024). doi:10.1007/978-1-0716-3969-6_2.

33. Dipalo, M. et al. Plasmonic meta-electrodes allow intracellular recordings at network level on high-density CMOS-multi-electrode arrays. Nat. Nanotechnol. 13, 965–971 (2018).

34. Schindelin, J., et al. Fiji: an open-source platform for biological-image analysis, Nat. Methods 9, 676–682 (2012).

35. Broadhead, M. J. et al. PSD95 nanoclusters are postsynaptic building blocks in hippocampus circuits. Sci. Rep. 6, 24626 (2016).

36. Blackburn, J. et al. Astrocyte regional heterogeneity revealed through machine learning-based glial neuroanatomical assays. J. Comp. Neurol. 529, 2464–2483 (2021).

37. Yang, B. et al. Neuron Image Segmentation via Learning Deep Features and Enhancing Weak Neuronal Structures. IEEE J. Biomed. Health Inform. 25, 1634–1645 (2021).

38. Reactive astrocytes secrete the chaperone HSPB1 to mediate neuroprotection | Science Advances. https://www.science.org/doi/10.1126/sciadv.adk9884.

39. Luijerink, L., Rodriguez, M. & Machaalani, R. Quantifying GFAP immunohistochemistry in the brain - Introduction of the Reactivity score (R-score) and how it compares to other methodologies. J. Neurosci. Methods 402, 110025 (2024).

40. Dahari, I., Weiss, O. E., Ayubi, A., Baranes, D. & Minnes, R. A method for quantifying parallel growth between neuronal dendritic branches in vitro. PLOS ONE 20, e0335919 (2025).

41. Leguey, I. et al. Patterns of Dendritic Basal Field Orientation of Pyramidal Neurons in the Rat Somatosensory Cortex. eNeuro 5, ENEURO.0142-18.2018 (2019).

42. Design and validation of a tool for neurite tracing and analysis in fluorescence microscopy images - Meijering - 2004 - Cytometry Part A - Wiley Online Library. https://onlinelibrary.wiley.com/doi/10.1002/cyto.a.20022.

