## Supplementary Information for "Hierarchical dendrite-inspired organic bioelectronic interfaces for neuronal integration"

### Supplementary Figure 1

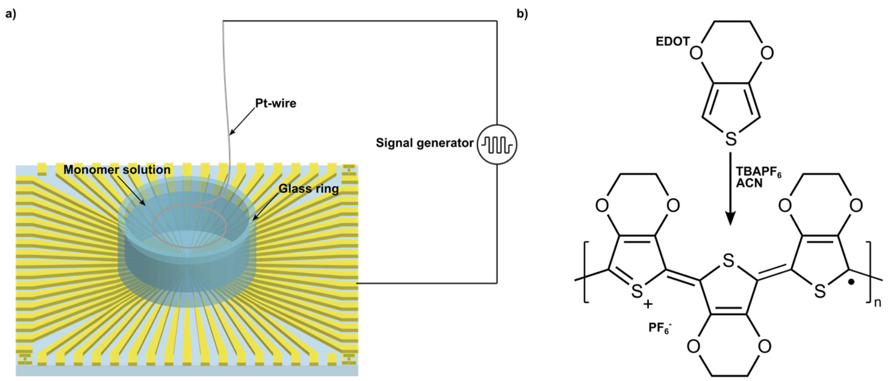


**Supplementary Figure 1 | Electrodeposition setup on MEAs. a**, Electrodeposition setup for the electrodeposition of PEDOT:$PF_6$ electrodes on MEAs. **b**, Chemical reaction of the PEDOT:$PF_6$ polymerization.

### Supplementary Figure 2

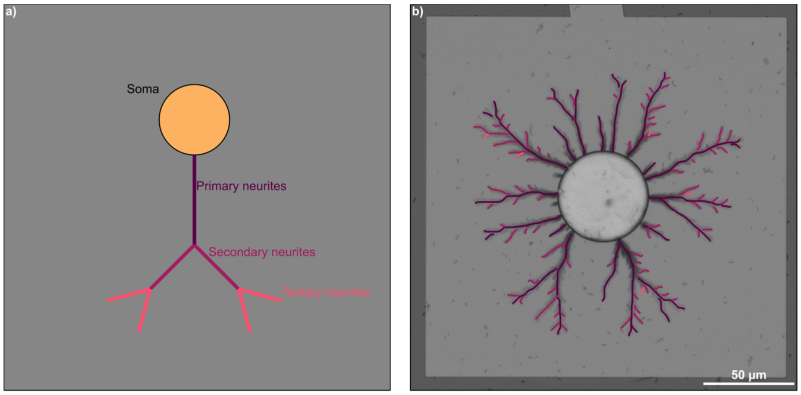


**Supplementary Figure 2 | Neuron inspired tracing of PEDOT:$PF_6$ fibers. a**, Neurite classification structure of biological neurons. **b**, Equivalent tracing on fiber electrodes following the typical neurite classification. Branches that split with equal length were classified as both primary branches.

### Supplementary Figure 3

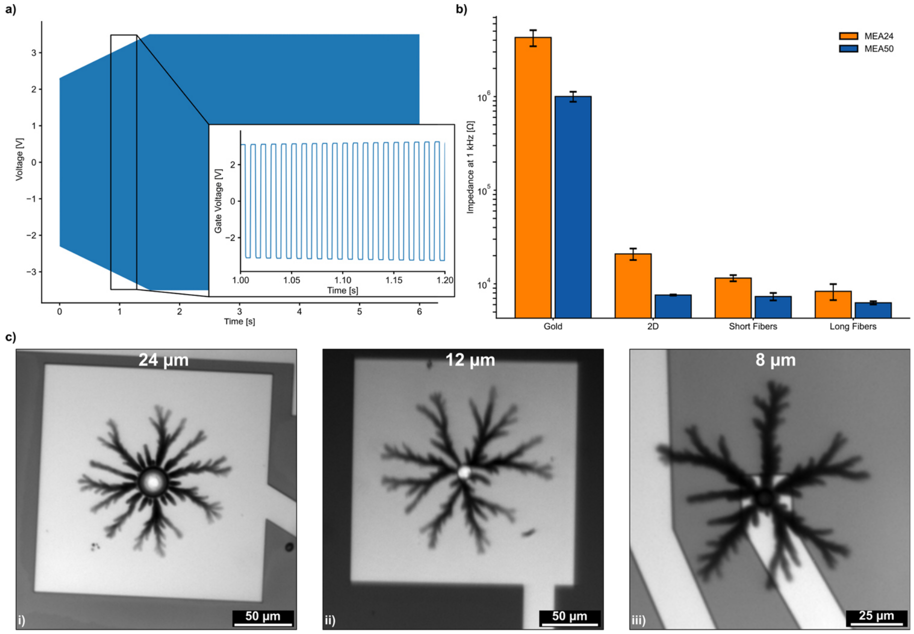


**Supplementary Figure 3 | Morphology preserving downscaling of the fiber growth. a**, example voltage trace applied for the scaled down electrodes, showing the stepwise increase during the initial deposition. **b**, Impedance at 1 kHz comparing 24 µm and 50 µm diameter electrodes. **c**, example optical images showing the fiber deposition scaled to 24 (**i**), 12 (**ii**) and 8 µm (**iii**) diameter electrodes.

To match the electrode opening with the size of neuronal somas, we chose an electrode opening of 24 µm. To this regard the fiber deposition was scaled down to achieve a similar structure independent from the electrode size. Applying the same signal to a smaller electrode leads to strong, film-like, deposition in the initial segment instead of the desired fiber growth. To prevent this initial strong deposition, the initial voltage is reduced in relation to the electrodes diameter and increased gradually until reaching 3.5 V (Supp. Figure 3a). The impedance at 1 kHz of the MEAs with a diameter of 50 µm and 24 µm was compared (Supp. Figure 3b), showing that the impedance values for the fiber electrodes are similar for both electrode diameters.

The starting voltage is set according to the following equation:

$$V_{start} = (3.5\,V - V_{oxidation}) * \frac{r_{new}}{50\ \mu m} + V_{oxidation}$$

Where $V_{oxidation}$ denotes the oxidation voltage of EDOT without a reference electrode (here 1.3 V) and $r_{new}$ denotes the new electrode diameter. The step size of the voltage increase was determined to be 8 mV per pulse. The long fiber depositions were scaled down to electrodes with a diameter of 24 µm, 12 µm and 8 µm by applying a starting voltage of 2.3 V, 1.75 V and 1.57 V, respectively. Example optical micrographs can be seen in Supp. Figure 3c, showing a similar structure throughout the different electrode openings.

### Supplementary Figure 4

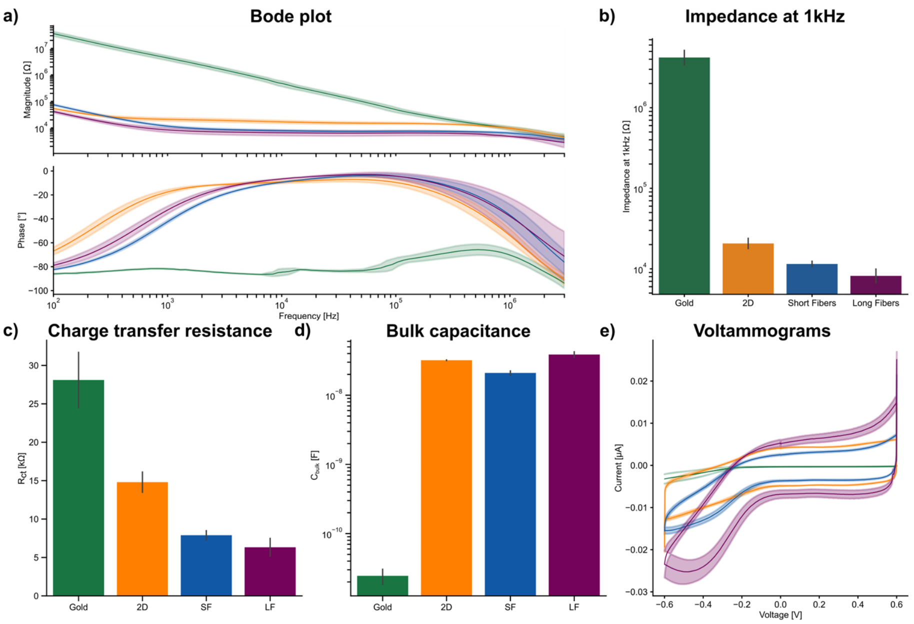


**Supplementary Figure 4 | Electrochemical characterization of gold, 2D, SF, and LF electrodes. a-d,** Impedance analysis on gold, 2D, SF and LF electrodes: showing the magnitude and phase (**a**), the impedance at 1 kHz (**b**), the extracted charge transfer resistance (**c**) and bulk capacitance (**d**).**c**, the cyclic voltammogram of the 2D, SF, LF and gold electrodes. Bode plots and voltammograms depict the mean ± sd as line and shadow, respectively. Bar plots depict the mean ± sd.

### Supplementary Figure 5

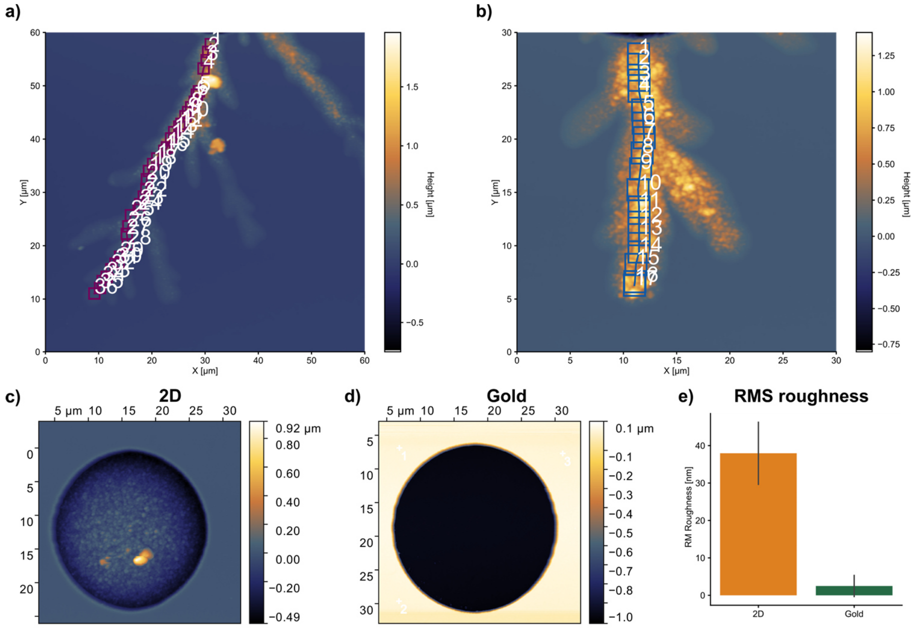


**Supplementary Figure 5 | Topography analysis on gold, 2D, SF and LF electrodes. a**,**b**, Example areas determined for the RMS roughness calculation on LF (**a**) and SF (**b**) branches. **c**, Representative AFM micrographs of 2D elctrode. **d**, Representative AFM micrograph of a gold electrode. **e**, RMS roughness. 2D and gold electrodes. Graphs represent the mean ± sd.

### Supplementary Figure 6

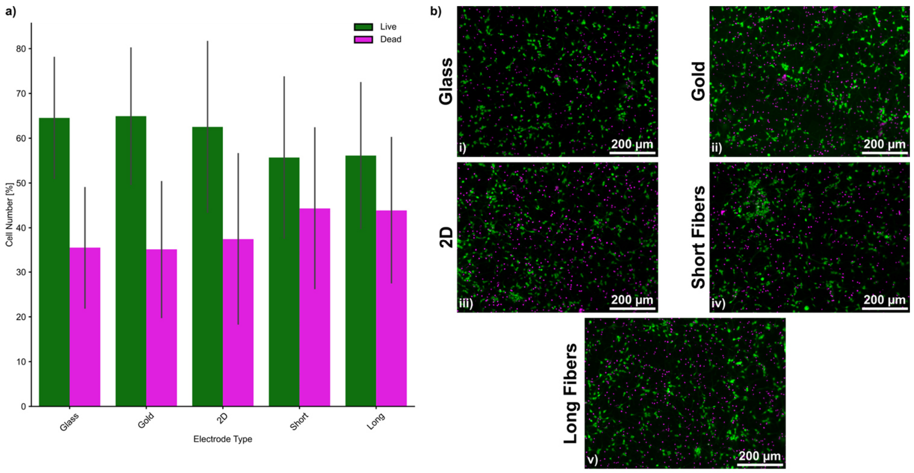


**Supplementary Figure 6 | Biocompatibility assay at DIV2. a**, number of live (green) and dead (dead) cells. **b**, representative images of the glass control (i) and MEAs with gold (ii), 2D (iii), SF (iv) and LF (v) electrodes showing live (CalceinAM) and dead (EtHD) cells. Brightness and contrast were linearly and identically adjusted for all micrographs. Graph represents the mean ± sd.

### Supplementary Figure 7

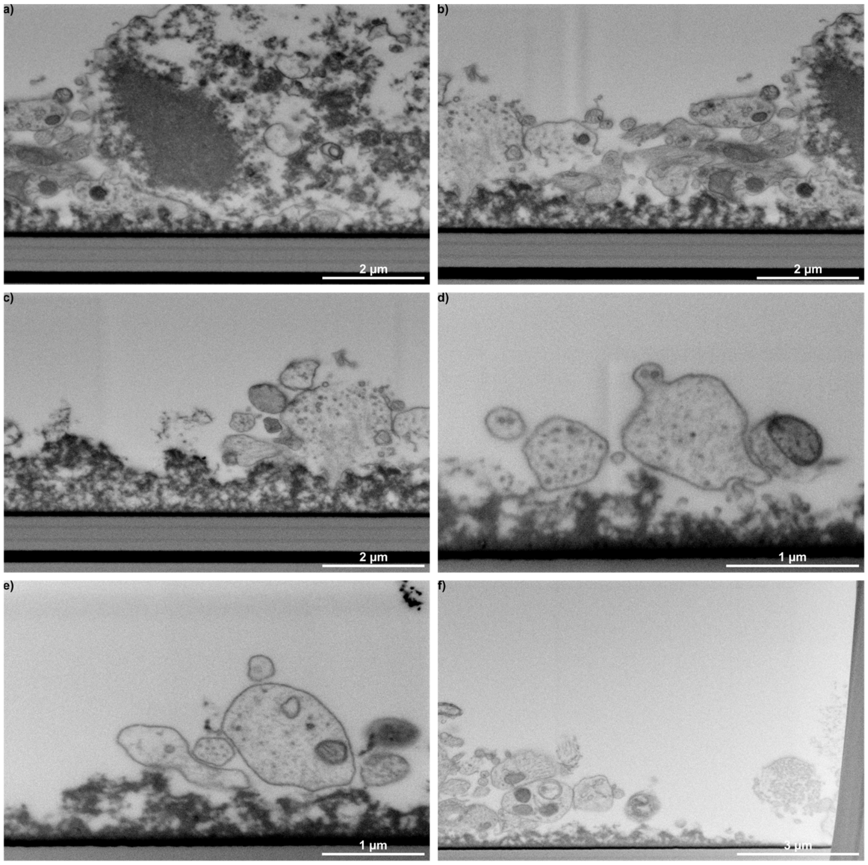


**Supplementary Figure 7 | FIB/SEM cross sections of neurites. a-f,** Example cross sections showing cross sections of neurites on top of the branch of a long fiber electrode.

### Supplementary Figure 8

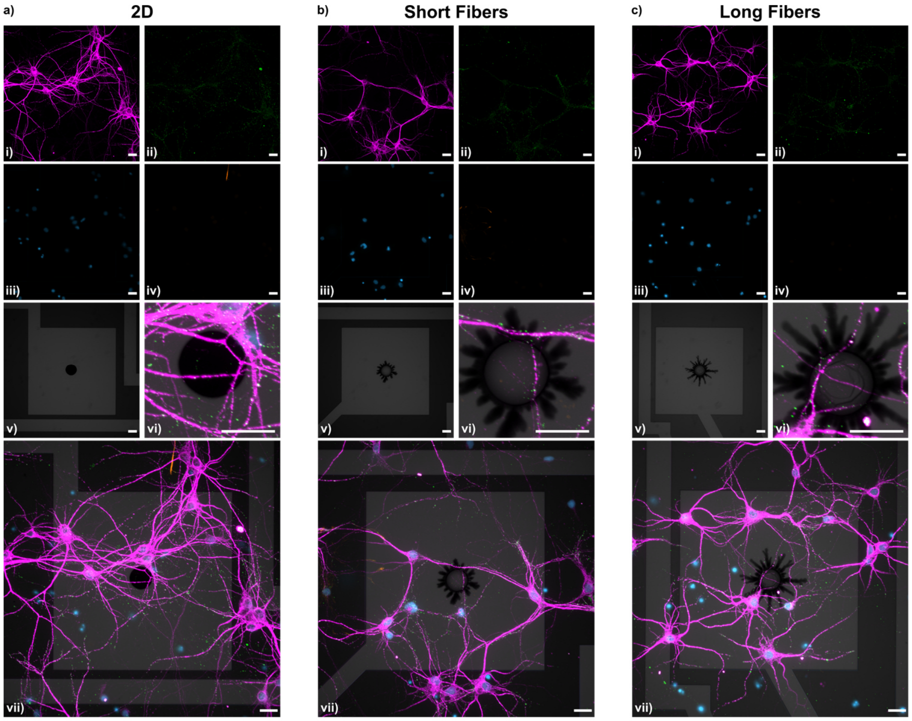


**Supplementary Figure 8 | Immunostainings on DIV10. a**-**c**, Fluorescence micrographs depicting dendrites (MAP2; **i**), excitatory postsynaptic sites (PSD95; **ii**), nuclei (DAPI; **iii**), astrocytes (GFAP; **iv**), electrode structure (brightfield; **v**) and the overlay in the electrode's ROI (**vi**) and full field of view (**vii**)for 2D (**a**), SF (**b**) and LF (**c**) electrodes on DIV10. Scalebars denote 20 µm. Brightness and contrast were linearly and identically adjusted for all micrographs.

### Supplementary Figure 9:

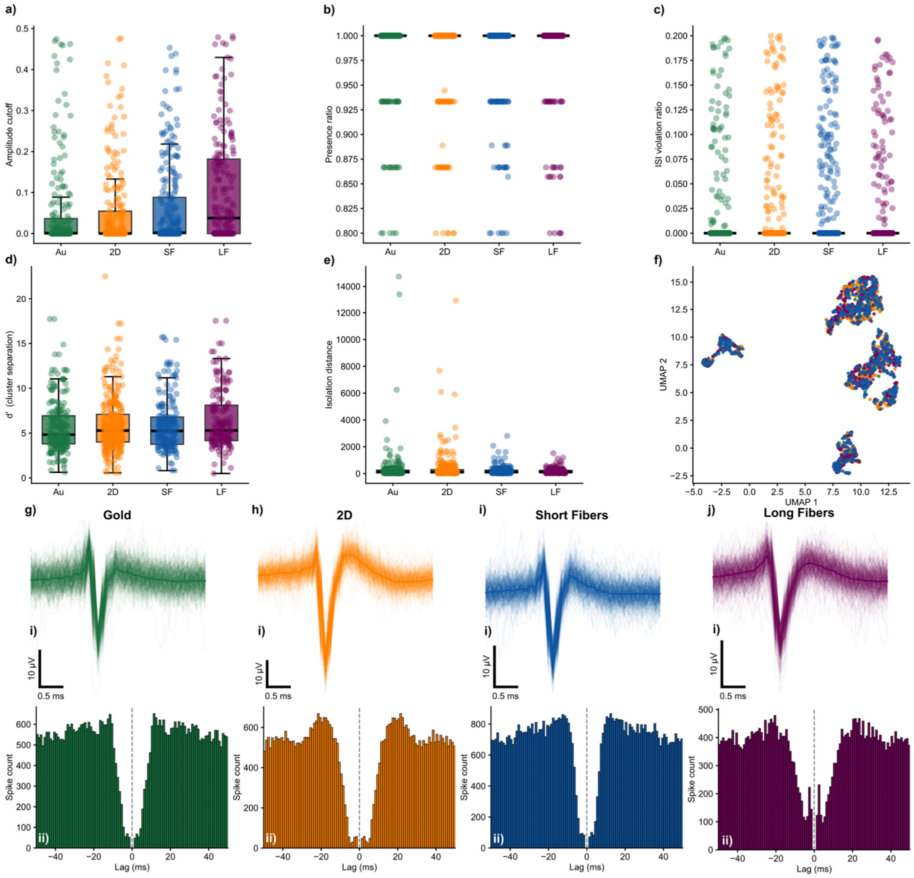


**Supplementary Figure 9 | Quality metrics of spike sorting. a-e**, Quality metrics extracted from the curated units, showing the amplitude cutoff (**a**), the presence ratio (**b**), the ISI violation ratio (**c**), d' prime (**d**) and isolation distance (**e**). **f**, Clustering units by electrode type, showing uniform distribution across all clusters. **g-j**, Representative units showing the overlayed waveforms (**i**) and autocorrelograms (**ii**) for gold (**g**), 2D (**h**), SF (**i**) and LF (**j**) electrodes

**Supplementary Figure 10:**

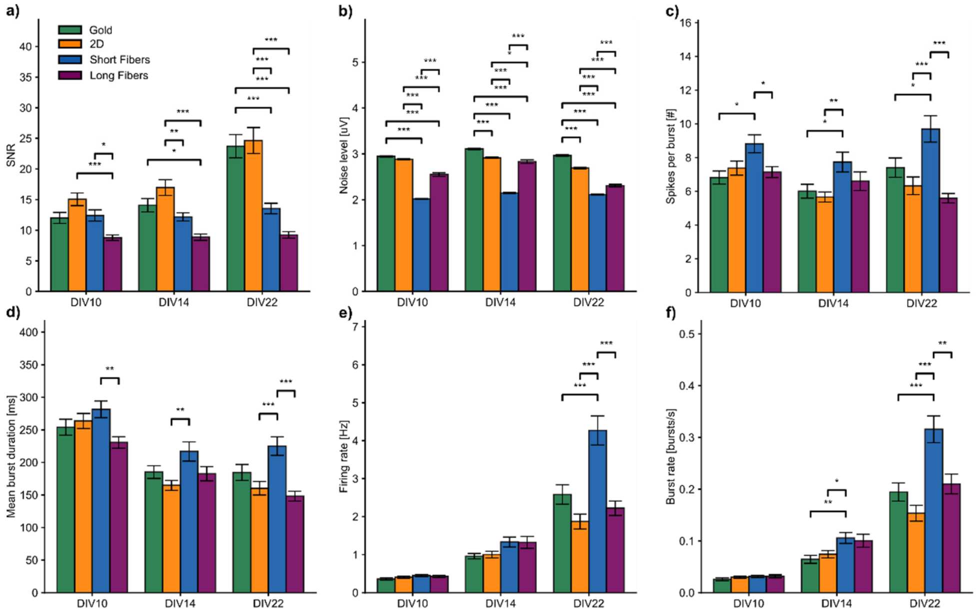


**Supplementary Figure 10 | Additional metrics of gold, 2D, SF and LF electrodes. a-f**, Additional extracted metrics for the gold, 2D, SF and LF electrodes showing the SNR (**a**), noise level (**b**), number of spikes per burst (**c**), mean burst duration (**d**), firing rate (**e**) and burst rate (**f**). Data represent n = 3 per condition from N = 3 independent experiments for each DIV. Graphs show the mean ± sem. Statistical significance was determined as previously described (* $p < 0.05$; ** $p < 0.01$, *** $p < 0.001$).

**Supplementary Figure 11:**

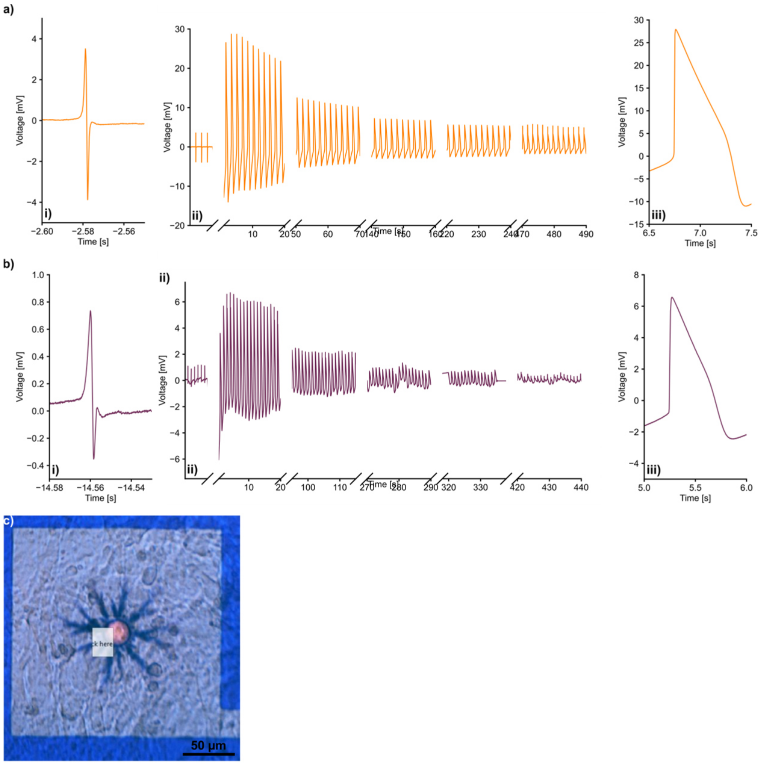


**Supplementary Figure 11 | Preliminary optoporations on hiPSC-derived cardiomyocytes. a**, optoporation on a 2D electrode, showing the extracellular field potential (**i**), mixed (**ii**) and intracellular action potential (**iii**) signals. **b**, optoporation on LF electrode, showing the extracellular field potential (**i**), mixed (**ii**) and intracellular action potential (**iii**) signals. **c**, example optical image of hiPSC-CMs cultured on a LF electrode. The top left corner of the square denotes the laser spot position for optoporation.

### Supplementary Figure 12

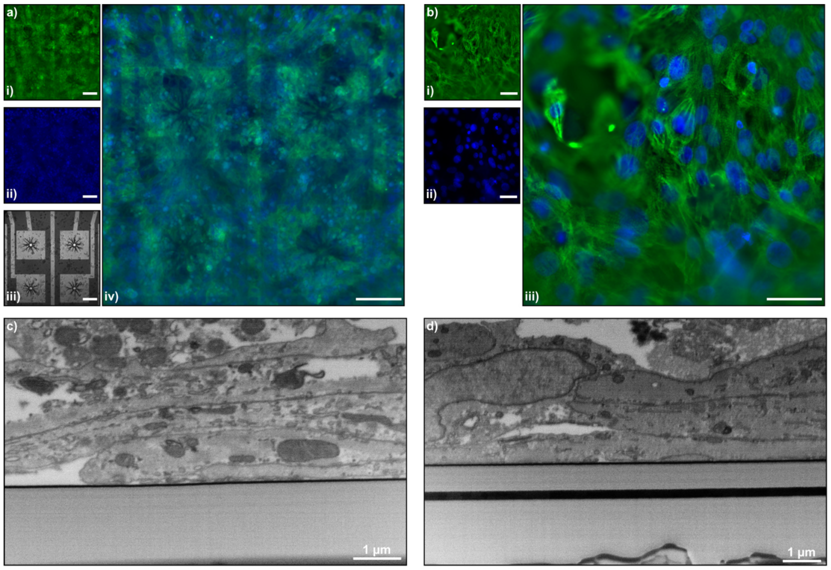


**Supplementary Figure 12 | Immunostaining and FIB/SEM cuts on hiPSC-CMs. a**,**b**, Immunostaining depicting the nuclei (DAPI; **i**), cardiac Troponin T (cTnT/TNNT2; **ii**) and electrode structure (brightfield; **iii**) of hiPSC-CMs at DIV14 showing on a LF electrode array (**a**) and ROI (**b**). **c**,**d**, FIB/SEM cross-sections of cardiomyocytes on the passivation layer of the MEA.

### Supplementary Figure 13:

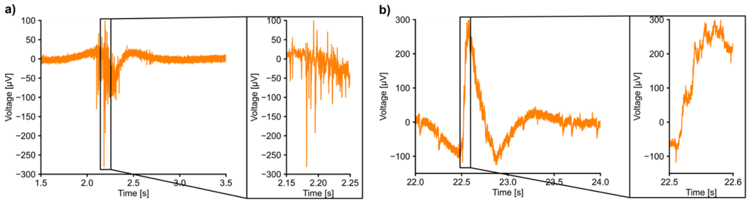


**Supplementary Figure 13 | Optoporation on a 2D electrode on Neurons. a**, Extracellular signal of extracellular action potentials before optoporation. **b**, in-cell signal after optporation.
